# Biologically Realistic Computational Primitives of Neocortex Implemented on Neuromorphic Hardware Improve Vision Transformer Performance

**DOI:** 10.1101/2024.10.06.616839

**Authors:** Asim Iqbal, Hassan Mahmood, Greg J. Stuart, Gord Fishell, Suraj Honnuraiah

**Author notes:** Corresponding authors: Asim Iqbal, Gord Fishell and Suraj Honnuraiah. Senior author and Lead contact (Suraj Honnuraiah).

## Abstract

Understanding the computational principles of the brain and replicating them on neuromorphic hardware and modern deep learning architectures is crucial for advancing neuro-inspired AI (NeuroAI). Here, we develop an experimentally-constrained biophysical network model of neocortical circuit motifs, focusing on layers 2-3 of the primary visual cortex (V1). We investigate the role of four major cortical interneuron classes in a competitive-cooperative computational primitive and validate these circuit motifs implemented soft winner-take-all (sWTA) computation for gain modulation, signal restoration, and context-dependent multistability. Using a novel parameter mapping technique, we configured IBM’s TrueNorth (TN) chip to implement sWTA computations, mirroring biological neural dynamics. Retrospectively, we observed a strong correspondence between the biophysical model and the TN hardware parameters, particularly in the roles of four key inhibitory neuron classes: Parvalbumin (feedforward inhibition), Somatostatin (feedback inhibition), VIP (disinhibition), and LAMP5 (gain normalization). Moreover, the sparse coupling of this sWTA motif was also able to simulate a two-state neural state machine on the TN chip, replicating working memory dynamics essential for cognitive tasks. Additionally, integrating the sWTA computation as a preprocessing layer in the Vision Transformer (ViT) enhanced its performance on the MNIST digit classification task, demonstrating improved generalization to previously unseen data and suggesting a mechanism akin to zero-shot learning. Our approach provides a framework for translating brain-inspired computations to neuromorphic hardware, with potential applications on platforms like Intel’s Loihi2 and IBM’s Northpole. By integrating biophysically accurate models with neuromorphic hardware and advanced machine learning techniques, we offer a comprehensive roadmap for embedding neural computation into NeuroAI systems.

## Introduction

Recent advances in machine learning and computational neuroscience have significantly accelerated the progress toward the development of synthetic cognitive agents with artificial general intelligence (AGI). Vision transformers (Dosovitskiy et al. [2020]) and natural language models (Shanahan et al. [2023]) have achieved notable success in image recognition and natural language processing. However, despite surpassing human performance in specific tasks like chess ([Campbell et al. 2002]) and Go (Silver et al. [2017]), AI systems still encounter significant challenges when learning in novel environments. These systems require substantially more computational resources and annotated data than biological brains.

This disparity may also arise from fundamental differences in how artificial and biological neural networks process information. In this work, we investigate the potential of reverse-engineering the brain’s computational principles and integrating them into AI systems. This exploration aligns with the core tenet of the NeuroAI approach (Zador et al. [2023]), aiming to bridge the existing gap between artificial and biological intelligence.

The execution of cognitive behavior in the brain relies on the ability to select actions based on external stimuli and context (Dayan [2008]). In animals, the learning of state-dependent sensorimotor mappings (Asaad et al. [2000], Banerjee et al. [2020], Xu et al. [2022], Condylis et al. [2020]) is primarily mediated by the neocortex, which facilitates cognition through computations enabled through its modular, laminar microcircuits. These microcircuits consist of excitatory and inhibitory neurons, including four major inhibitory classes — parvalbumin (PV), somatostatin (SST), vasoactive intestinal peptide (VIP), and Lamp5 (Rudy et al. [2011], [Tremblay et al. 2016]). These interneurons play a crucial role in regulating state-dependent computations, performing tasks such as arithmetic, logical operations, timing, and gain modulation (Fishell and Kepecs [2020], Kepecs and Fishell [2014], Ferguson and Cardin [2020], Niell and Scanziani [2021]). Importantly, these four inhibitory neuron classes are conserved across cortical regions and species (Pfeffer et al. [2013a], Campagnola et al. [2022]), indicating that their computational logic is generalizable for diverse high-order tasks, including motor execution and working memory.

Several key candidate computational principles have been proposed to elucidate neocortical function, including normalization (Carandini and Heeger [2012]), dynamic field theory (Schöner and Spencer [2016]), attractor networks (Vyas et al. [2020]), predictive coding (Keller and Mrsic-Flogel [2018]), Bayesian inference (Bastos et al. [2012]) and winner-take-all (WTA) computations (Douglas and Martin [2007]). However, direct evidence at the level of microcircuit and biological hardware implementation remains limited. Among these, the WTA mechanism is amenable to neocortical architecture and combines key elements of these various computational approaches. By employing competitive-cooperative dynamics, the WTA mechanism facilitates selective amplification and noise minimization, thus enhancing signal restoration ([Douglas and Martin 2007]). These characteristics resemble signal processing in both primary sensory and motor cortices, where superficial pyramidal neurons receive sparse and weak thalamic inputs that require amplification to extract relevant information (Balcioglu et al. [2023], Lien and Scanziani [2018], Bopp et al. [2017], Binzegger et al. [2004]). Consequently, the WTA mechanism may be a fundamental computational strategy employed by cortical circuits and represent a ubiquitous computational strategy implemented by the neocortex.

In addition, WTA models show considerable promise for neuromorphic hardware (Mead, 1990; 2023), especially in energy-efficient, real-time processing (Chicca et al. [2014], Qiao et al. [2015], Indiveri and Sandamirskaya [2019]). To execute such computations in silico, IBM’s TrueNorth (TN) chip offers a tractable platform for integrating brain-inspired principles. For example, it features a reconfigurable, asynchronous, multi-core digital architecture optimized for real-time, ultra-low-power, event-driven processing with physical neurons (Modha et al. 2023a] Merolla et al. 2014], [Neckar et al. [2018a]). As a result, TN is especially well-suited for implementing brain-like computations (Indiveri and Sandamirskaya [2019]). While prior research has focused on leveraging statistical relationships among neuronal populations to emulate biological circuits on TrueNorth hardware (Imam [2021]), developing generalizable techniques for integrating diverse biophysical and theoretical models remains an open challenge. In this work, we demonstrate that by employing biophysically realistic computational principles, parameters of IBM TrueNorth - such as thresholds, leak rates, and crossbar weights, can be modeled to reflect those found in cortical microcircuits. By utilizing this approach, TrueNorth (TN) hardware can be programmed to perform computations analogous to those observed in simplified, biologically realistic V1 cortical circuit motifs, that may potentially underlie key V1 functions such as orientation and direction tuning (Rossi et al. [2020], Niell and Stryker [2008], Douglas and Martin [2007], Hubel and Wiesel [1962].

Our goal here was not to build an exhaustive model of V1, such as that outlined in Billeh et al. [2020], but rather to design a simplified, generalizable circuit motif that validates the core computational principles utilized in cortical processing. The retrospective analysis confirmed that the optimal parameters for configuring TN hardware to display sWTA dynamics closely aligned with the primary functions of different interneuron classes. Furthermore, our findings demonstrated that hardware-optimized abstractions could effectively replicate biological circuits. Finally, to test the functionality of this approach, we investigated whether integrating this hardware-constrained sWTA computation could be utilized to implement a neural state machine for working memory or to enhance the performance of state-of-the-art deep learning models such as Vision Transformers (ViT). For the former, we successfully achieved persistent activity in the TN hardware by leveraging sparsely coupled sWTA motifs, a critical requirement for instituting working memory. Additionally, when this approach was applied as a pre-processing layer into the ViT architecture, we observed a substantial increase in classification accuracy for previously unseen test data. Together these results suggest that adapting biophysical principles to neuromorphic chips may offer a promising pathway for NeuroAI performance.

## Results

Our objective is to extract general principles of neocortical function, such as soft Winner-Take-All (sWTA), and implement them efficiently on neuromorphic hardware. By leveraging the hardware’s parametric constraints, we aim to apply these simplified neocortical computations to enhance working memory capabilities. This approach not only mimics brain-like processing but also has the potential to improve AI models’ performance across various machine learning tasks, bridging the gap between neuroscience and artificial intelligence.

### Biophysical model implementation of neocortical circuit motifs

Sensory information processing in primary sensory cortices, such as V1 relies on pyramidal neurons integrating bottom-up signals with top-down feedback from higher-order visual areas. Key components in this process include recurrent excitation, feedforward and feedback inhibition, disinhibition, and divisive normalization. These functions are primarily mediated by parvalbumin (PV), somatostatin (SST), vasoactive intestinal peptide (VIP), and lysosomal-associated membrane protein 5 (LAMP5) interneurons, respectively, working together to selectively amplify thalamic inputs (Reinhold et al. [2015], Reinhold et al. [2015], Pfeffer et al. [2013a], Lien and Scanziani [2013]). Mouse V1 exhibits strong recurrent connections among layer 2/3 (L2/3) pyramidal neurons (Ko et al. [2011], Harris and Mrsic-Flogel [2013], Rossi et al. [2020]), with PV interneurons providing local feedforward inhibition and SST interneurons delivering global feedback inhibition targeting axon initial segments and L2/3 dendrites (Schneider-Mizell et al. [2021], Atallah et al. [2012], Naka et al. [2019], Adesnik and Scanziani [2010]). VIP interneurons modulate feedback inhibition (Karnani et al. [2016], Pfeffer et al. [2013a]), while LAMP5 cells regulate top-down and bottom-up signals through normalization (Ibrahim et al. [2021], Huang et al. [2023], Malina

Consistent with earlier studies, our experimental data confirmed that L2/3 pyramidal neurons receive strong inhibition during bottom-up sensory input stimulation in V1 (**Figure 1A-C**). To dissect the specific contributions of PV and SST interneurons, we employed optogenetics in PV-Cre and SST-Cre mice and analyzed how their activation modulated the current-frequency (f/I) response of L2/3 pyramidal neurons (**Figure 1D-F**). Additionally, top-down auditory cortex inputs, which primarily convey contextual information, were found to target Lamp5 expressing neurogliaform interneurons (**Figure 1G-J**). Though not explicitly tested, we modeled Lamp5-mediated global inhibition via volumetric transmission (Ibrahim et al. [2021], Huang et al. [2023]) affecting L2/3 pyramidal neurons. Finally, VIP interneurons, although not directly included, likely modulated SST-mediated inhibition through disinhibition (Karnani et al. [2016], Pfeffer et al. [2013a]).

**Figure 1:**
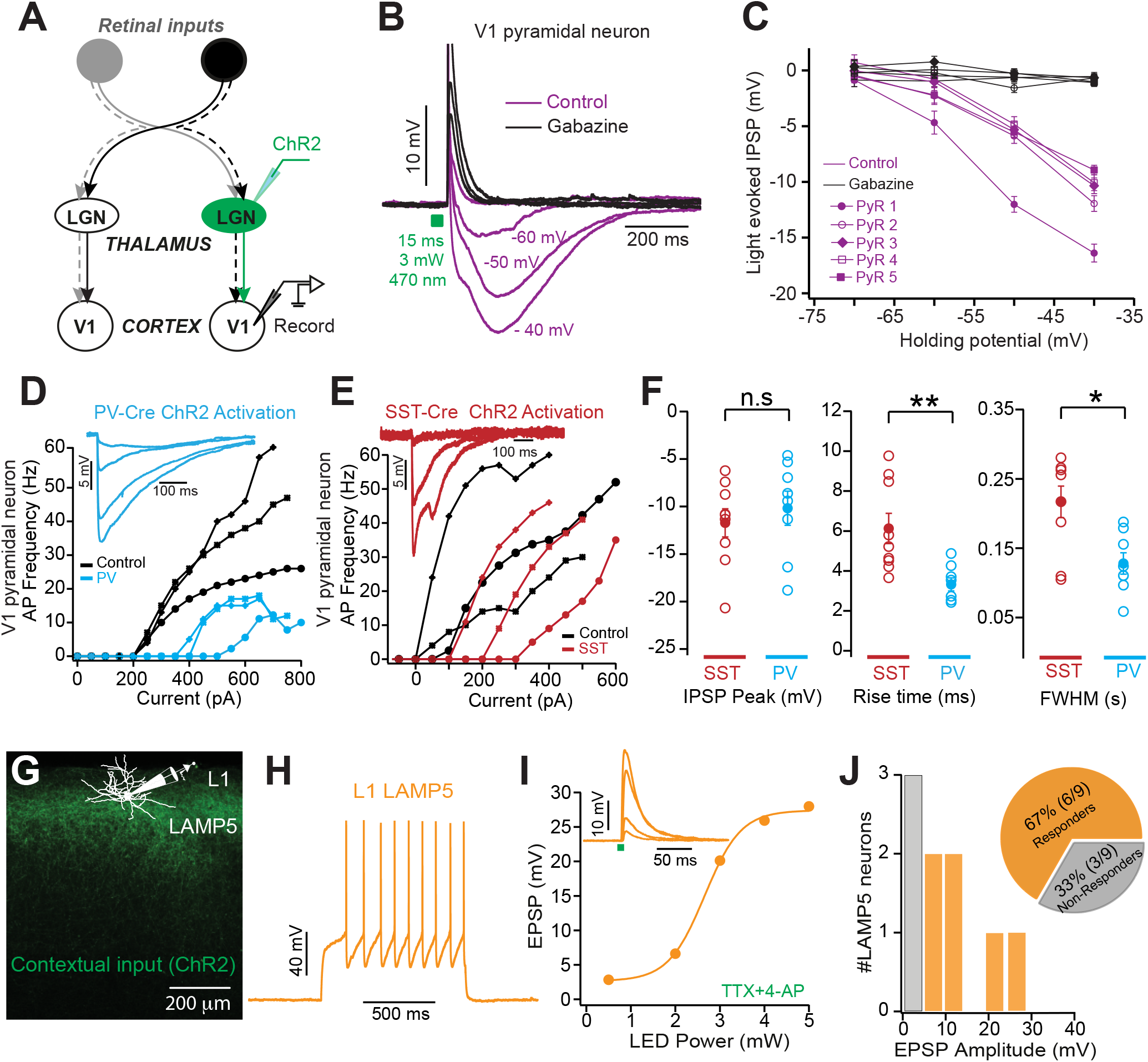
A) Schematic of the experimental arrangement showing ChR2 injections in LGN, and recordings in pyramidal neuron in V1. B) Average light-evoked synaptic responses (10 trails) in a representative layer 2/3 pyramidal neuron in V1 at the indicated holding potentials in control (magenta) and in the presence of GABAzine (black). C) Summary data showing IPSP amplitude versus holding potential in control (magenta) and in the presence of GABAzine (black) in different binocular layer 2/3 pyramidal neurons. D-E) Impact of PV and SST neuron activation on the f/I curves of L2/3 pyramidal neurons. Voltage responses of binocular pyramidal neurons at different resting membrane potentials during cre-dependent activation of PV (D) and SST (E) inhibitory neurons. F) Quantification of peak hyperpolarization (left panel), rise time (middle panel), and full-width half maximum (right panel) of the inhibition evoked by SOM and PV neuron activation in binocular pyramidal neurons at a resting membrane potential of -45 mV. G) Schematic showing contextual inputs from the Auditory cortex to layer 1 Lamp5 neurons in the somatosensory cortex (S1). H) Suprathreshold voltage response of an L1 Lamp5 interneuron to a somatic current injection of 200 pA. I) Amplitude of EPSPs in a layer 1 neuron in S1 versus LED power (470 nm, 2 ms) in the presence of TTX+4-AP. Inset: Example EPSPs during photo-activation of auditory cortex axons with increasing power (0.24 to 1 mW). J) Histogram of EPSP amplitude in layer 1 interneurons receiving (cyan) and not receiving (grey) A1 input. Inset: Pie chart of the distribution of responding (cyan) and non-responding (grey) layer 1 interneurons.

Next, we built a biophysically detailed network model in the NEURON simulation environment to validate whether the simplified cortical circuit motifs obtained from V1 indeed implemented sWTA computations. To do this, we incorporated excitatory and inhibitory cell types with diverse spiking patterns in a conductance-based Hodgkin-Huxley network model (**Supplementary Fig. 2A-D**). Synaptic parameters were constrained by our experimental data (**Fig. 1F**). Poisson-modulated excitatory synaptic inputs were used to assess the input-output (IO) function of L2/3 pyramidal neurons under PV and SST inhibition (**Supplementary Fig. 2A**). Synaptic weights were tuned to match experimentally observed inhibitory postsynaptic potentials, adjusting the pyramidal neuron f/I curve (**Supplementary Fig. 2D-F**). SST inhibition primarily influenced the slope of the IO function, while PV inhibition altered the offset (**Supplementary Fig. 2E-F**). Lamp5-mediated volumetric inhibition was represented as non-specific inhibition across pyramidal neuron dendrites, achieved by reducing both SST and recurrent excitatory weights. Notably, although VIP inhibition was not explicitly modeled, its effects were captured by reducing SST weights.

Based on experimental data, we developed a generalized cortical microcircuit model comprising 10 pyramidal neurons, 10 PV interneurons, 1 SST interneuron, and 1 Lamp5 interneuron, with their biophysical properties constrained by our mouse V1 physiology data (**Figure 2A**). We examined the computational behavior of this model under Poisson-modulated excitatory synaptic inputs mimicking thalamic activity, where stronger inputs were selectively amplified, and weaker inputs suppressed (**Figure 2B**). Stronger thalamic inputs represent a tuned orientation or direction information carried to L2/3 pyramidal neurons in V1. The soft winner-take-all (sWTA) mechanism enables neural circuits to prioritize the strongest input by balancing competitive and cooperative interactions. Non-linear amplification of stronger inputs, coupled with suppression of weaker ones, forms the core of sWTA computations. Consistent with previous findings (Douglas et al. [1995], Somers et al. [1995], Reinhold et al. [2015]), our model reproduces through recurrent excitation and lateral feedback inhibition (**Figure 2B-C**). Using this model, we assessed how various interneuron populations contribute to sWTA computations, focusing on their role in enhancing responses to strongly stimulated neurons (Pyr 4) while suppressing weaker responses (Pyr 1-3, 5-7; **Figure 2C**). By modulating the conductances of PV, SST, VIP, and Lamp5 interneurons within physiological ranges, we quantified their influence on pyramidal neuron dynamics (**Figure 2D-E**). PV and SST inhibition independently shaped the width and gain of the sWTA function, while Lamp5-mediated inhibition primarily adjusted gain (**Figure 2F**). VIP-mediated disinhibition was examined by reducing SST synaptic weights.

**Figure 2:**
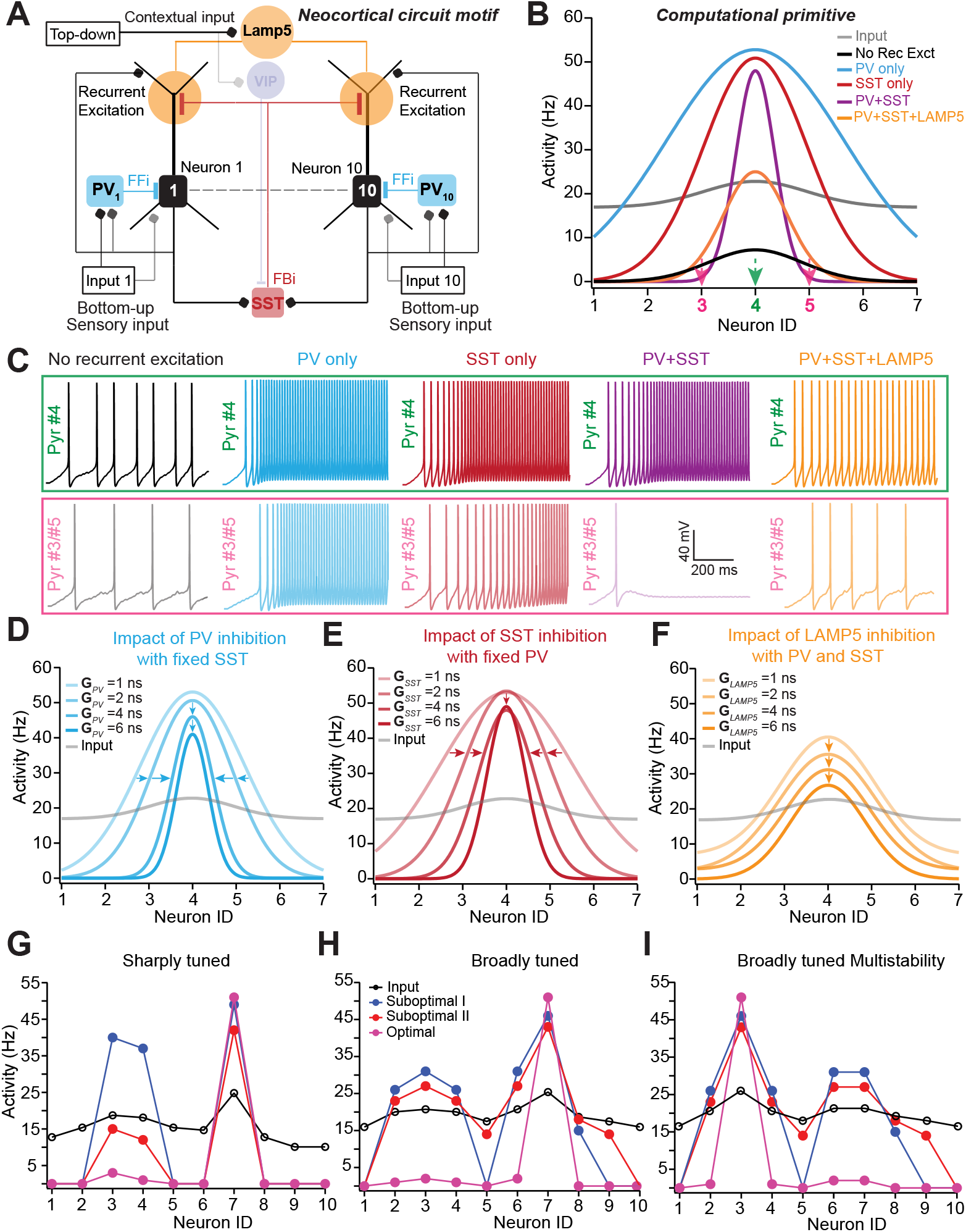
Biophysical implementation of experimentally validated neocortical circuit motifs using a conductance-based neural network model. A) Canonical neocortical circuit motif implemented using a conductance-based Hodgkin-Huxley neuron network model consisting of 10 pyramidal neurons, 10 PV interneurons providing feedforward and local feedback inhibition, and one common inhibitory neuron by LAMP5 for contextual modulation and one SST interneuron providing lateral inhibition. Sensory input is Poisson modulated excitatory synaptic inputs with varying frequencies. B) Computational primitives implemented by the neocortical circuit motifs performing non-linear input amplification, selective suppression, and soft-winner take-all (sWTA) computation. The contribution of various circuit elements like distinct interneurons: PV, SST, and LAMP5 mediated inhibition and recurrent excitation to sWTA computation is shown in various colors. Gaussian fits of the data points are used to extract quantifiable parameters such as mean and width. C) Voltage traces of representative pyramidal neurons receiving the highest (in green) and next to highest (in pink) input shown during various conditions: without recurrent excitation (black), with recurrent excitation and PV plus SST inhibition (magenta) along with LAMP5 inhibition (orange), and only SST (red) and only PV inhibition (cyan). D-F) Impact of modulating distinct inhibition on the computational primitives implemented by the circuit motifs in (A). Varying PV inhibitory conductance from 1-6 nS with a fixed SST inhibition of 1.5 nS along with recurrent excitation (D). Varying SST inhibitory conductance from 1-6 nS with a fixed PV inhibition of 1.5 nS along with recurrent excitation (E). Varying LAMP5 inhibitory conductance from 1-6 nS with an SST and PV inhibition along with recurrent excitation (F). G-I) Verification of sWTA properties such as nonlinear amplification and signal restoration for sharply and broadly (H) tuned inputs. Multistability and signal invariance for broadly (I) tuned inputs for suboptimal (blue, red) and optimal (magenta) parameters are derived from our proposed method.

We computed a selectivity index to evaluate the network’s ability to suppress weaker inputs while amplifying stronger ones. This index, calculated as the difference in pyramidal neuron responses to closely tuned inputs, revealed that recurrent excitation, along with PV and SST inhibition, is crucial for maintaining high selectivity (Supplementary Fig. 2G-I). Lamp5 inhibition modulates gain, preserving tuning specificity while enabling flexible responses to factors like attention and locomotion (Ferguson and Cardin [2020], Bugeon et al. [2022]).

Our model replicates key features of cortical circuits, showing both linear gain scaling and non-linear selectivity. These dynamics are governed by the balance between excitatory and inhibitory synaptic weights and the feedback inhibition threshold, modulated by network activity. Deviations from optimal weights diminished sparsity by amplifying secondary inputs (**Figure 2G-I**). Linear gain modulation enhanced weak thalamic inputs ((Oldenburg et al. [2024], Sievers et al. [2024], Lien and Scanziani [2018], Lien and Scanziani [2013]),), while non-linear computations such as signal restoration for sharply tuned, noise-embedded inputs (**Figure 2G-H**) — aligned with *in vivo* observations. Furthermore, the model captures hysteresis and multistability, enabling the circuit to amplify relevant inputs based on initial conditions or contextual cues (**Figure 2I**).

In summary, our model provides a biophysical foundation for a simplified and generalized computational mechanism, such as soft Winner-Take-All (sWTA), which may represent a universal computation in sensory cortices (Douglas and Martin [2007], Niell and Stryker [2008]), that might contribute to orientation and direction tuning in the visual cortex (Hubel and Wiesel [1962]), angular whisker tuning in the barrel cortex (Lavzin et al. [2012]), and frequency tuning in the auditory cortex (Kato et al. [2017]).

### Mapping neocortical algorithms onto IBM TrueNorth neuromorphic hardware

A few years ago, IBM released their neuromorphic TrueNorth (TN) chip, offering a reconfigurable, asynchronous, multi-core digital architecture ideal for implementing brain-inspired computations. We aimed to program the TN chip to implement a simplified sWTA computational primitive, inspired by the neocortex. A key challenge was that TN’s neural dynamics were governed by parameters such as thresholds, leaks, and crossbar weight that did not directly align with biophysical or artificial neural network models. Notably, these strongly resemble gain modulation regulated by the four interneuron types considered in the biophysical modeling described previously.

To implement an sWTA computation, we developed an automated gain-matching technique to match TN network dynamics to the biophysical model, enabling accurate parameter mapping (Appendix A1). Initially, we created an abstract rate-based model that mimicked the input-output (IO) function of the biophysical neurons. We then derived constraints to map these dynamics onto the TN hardware, producing a linear threshold response that closely approximated the physiological behavior of cortical neurons (**Figure 3A-C**). Inputs to the TN network were generated by configuring the on-chip neurons within neurosynaptic cores to produce a range of frequencies (**Figure 3D-E**). This allowed us to match the TN network’s IO gain to the abstract model (**Figure 3F**). Using contraction theory (Rutishauser and Douglas [2009a]; Appendix A3), we derived optimal TN parameters, enabling us to implement all sWTA operations, as we performed in our biophysical analysis of V1 processing, including signal restoration, hysteresis, and multi-stability (**Figure 3G-I**).

**Figure 3:**
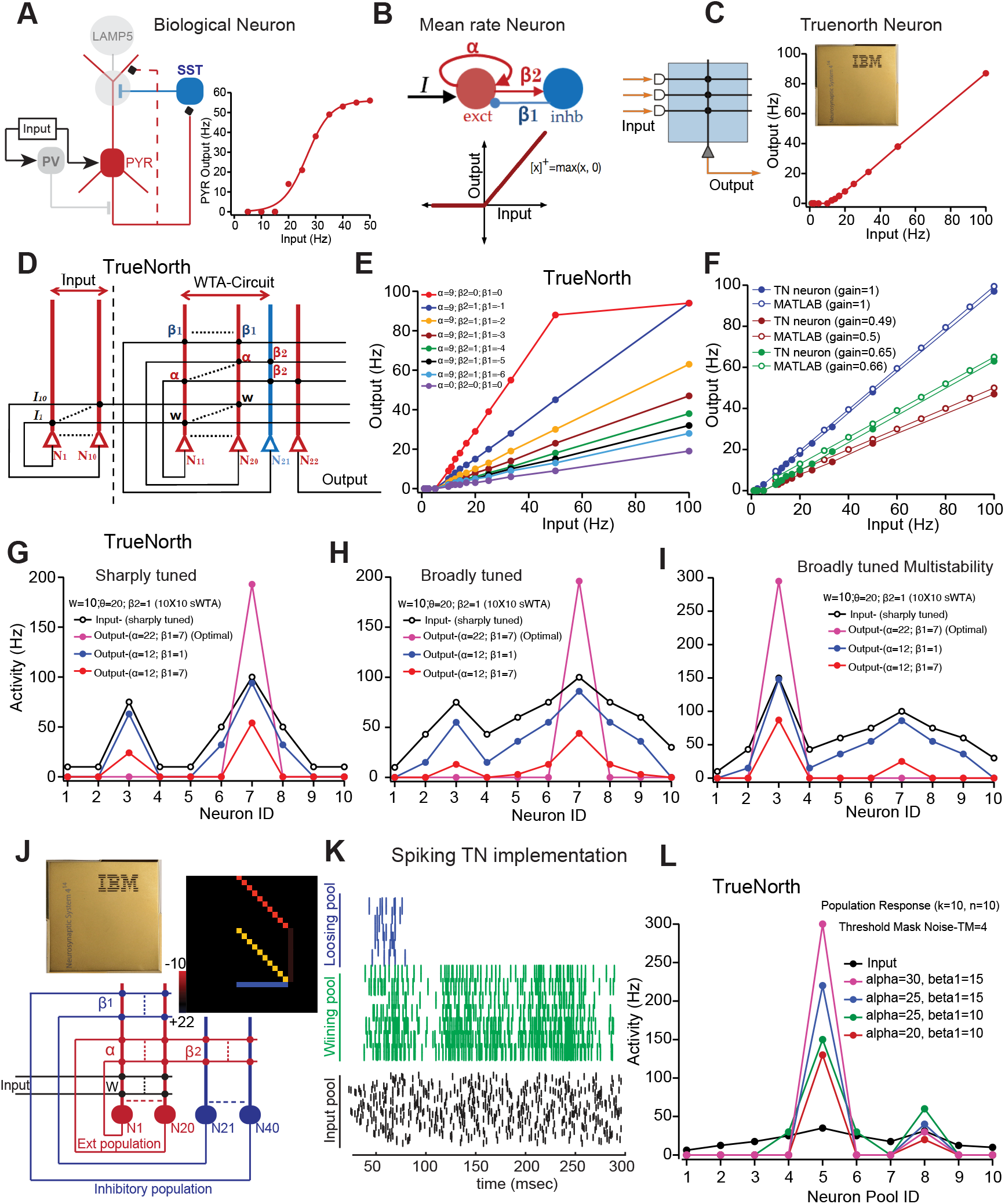
Configuring IBM TN neuromorphic hardware to implement neocortical circuit motifs and computational primitives. A) Left: Simplified biophysical circuit implementation (excitatory neurons in red; lateral inhibition in blue). Right: Transfer function of the L2/3 pyramidal neuron when stimulated with Poisson input. Stimulus frequency is varied between 5 to 50 Hz (in steps of 5 Hz) and the corresponding output response is plotted. B) Top: Abstract rate-encoding neuron model containing one excitatory (red) and inhibitory (blue) neuron with linear-threshold activation function (bottom). C) Left: Single TrueNorth neuron with crossbar weights along with input and output. Right: Transfer function of the TrueNorth neuron in a rate-encoding configuration similar to the transfer function of biological and abstract neuron models. D) Wiring diagram showing the connection configuration between 10 excitatory and 1 feedback inhibitory truenorth neurons for implementing the circuit motif shown in Figure 2A. E) Impact of recurrent excitation and feedback inhibition on the slope of the transfer function of the TrueNorth neuron shown in. F) Matching the slope/gain of the transfer function of the TN neuron with the abstract rate encoding neuron model implemented in software by tuning the TN parameters as described in Appendix A2, A3. G-I) Parameter tuning and verification of sWTA properties like nonlinear amplification and signal restoration in TN simulation and TN hardware to sharply tuned inputs (G), signal invariance property of sWTA to broadly tuned inputs (H), and multistability (I). Output response for optimal parameters is shown in magenta, inputs are shown in black, and responses for suboptimal parameters are shown in blue and red. J-L) Population-level implementation of sWTA in the spiking-mode configuration of Truenorth neural network.

After programming the TN chip to perform sWTA computations, we retrospectively compared TN parameters such as thresholds and leaks with those in our biophysical model. The optimized TN parameters closely aligned with the functions of the excitatory-inhibitory balance observed in the biophysical model (Supplementary Fig. 2E-F). Specifically, TN parameters such as threshold and leak mirrored the roles of PV and Lamp5-mediated inhibition in the biophysical model (Supplementary Fig. 3A-C). Moreover, the TN neurons replicated the recurrent excitation and global feedback inhibition motifs found in cortical circuits, mirroring the effects of SST interneurons on pyramidal neuron IO functions, as well as the role of VIP interneurons in disinhibiting this population (**Figure 3E**). Notably, the parameters derived using our gain-matching technique closely resembled those observed in experimentally constrained models of neocortical circuits across all conditions, highlighting the fidelity of TN hardware in simulating cortical computations (Supplementary Fig. 3D-F).

While our initial efforts efficiently mapped the sWTA computations in rate mode, we extended the method to implement population-level sWTA networks in spiking mode configuration (**Figure 3J**). This configuration, tested on a population of 10 excitatory neurons gated by shared inhibition, allowed us to translate rate-based computations into spiking dynamics, better reflecting neocortical organization. Under optimal conditions, the TN network’s dynamic range during sWTA closely matched the firing rates of cortical neurons in V1 (**Figure 3K-L**). We evaluated whether the TN network in spiking mode could implement sWTA computations under noisy conditions, simulating biological variability (**Figure 3K-L**). The noise was controlled using a parameter called threshold mask noise (TMN), which emulated spontaneous cortical activity. At TMN values up to 10, synaptic and spiking variability resulted in stable network dynamics that supported sWTA operations (Supplementary Fig. 3G-H). This demonstrated that TN spiking networks remain stable in noisy environments and avoid synchrony driven by inhibition. Without inhibition, excitatory activity would increase exponentially; however, when inhibitory neurons are activated, their gain is tuned to stabilize excitatory activity. This balance between positive and negative feedback forms an attractor state.

We next aimed to demonstrate a practical use case for sWTA circuit motifs in hardware applications, specifically by implementing a neural state machine (NSM). We hypothesized that NSMs could serve as foundational elements for complex cognitive tasks in robotics, and will benefit from the energy-efficient framework of sWTA networks. A key aspect of cognitive function is working memory, which allows for the retention of cue information even in the absence of stimuli, enabling appropriate action selection based on environmental cues. NSMs with working memory encode stimuli as distinct states, transitioning between them to support context-dependent tasks (e.g., Fuster and Alexander [1971], Wang [2001], Harvey et al. [2012]). Previous studies have shown that sparsely coupled sWTA motifs can sustain persistent activity (Neftci et al. [2013a], Rutishauser and Douglas [2009a]). Given the conserved use of circuit motifs across cortical areas, we hypothesized that this sWTA architecture would efficiently support working memory. To implement stable attractor states, TN hardware parameters were tuned to balance positive and negative feedback as outlined in Neftci et al. [2013a] to avoid inhibition-mediated synchrony, which is critical for maintaining persistent activity in an NSM using spiking dynamics. We tested whether these motifs could facilitate action selection in response to environmental cues while retaining information in their absence. Our results showed that TN neurosynaptic cores, initially designed for sensory sWTA, could be effectively repurposed for NSM implementation. Using sparsely coupled sWTA motifs, we achieved persistent activity in TN hardware, with time constants that aligned closely with experimental data.

We then evaluated the ability of this architecture to implement a two-state NSM. Transitions between states S1 and S2 were driven by input signals (X, Y) and regulated by pointer neurons (P12, P21) within the sparsely coupled sWTA motif (Supplementary Figure 4A-C). This configuration generated stable sWTA dynamics and persistent attractor states (Neftci et al. [2013a]; Appendix A3). Consistent with previous findings, gamma coupling through bidirectional excitatory weights in TN hardware sustained persistent activity even without external input. When noise was introduced, it disrupted synchronous firing but optimizing gamma coupling maintained stable persistence (Supplementary Fig. 4D). By fine-tuning the coupling strength, we identified the minimal gamma required for maintaining persistent activity across varying noise levels. Striking this balance was critical for stability, especially when environmental cues were unreliable. We further confirmed the stability of attractor states by removing one transition input, demonstrating that the circuit continued to sustain activity (Supplementary Fig. 4E).

In summary, we developed a two-state NSM where transitions were governed by input signals and the current state (Supplementary Fig. 4F). The sWTA dynamics facilitated smooth state transitions within a finite state automaton (FSA) framework. Efficiency analysis revealed that both time and energy in TN hardware scaled linearly with the number of states and computational load, contrasting with the quadratic scaling observed in Compass simulations. Notably, runtime on TN hardware was independent of firing rates, synapse activity, and neuron counts, with asynchronous state updates (Supplementary Fig. 4G-H; Appendix A2). This underscores the efficiency of neuromorphic hardware in implementing NSMs, supporting higher cognitive functions.

### Neocortex-inspired WTA implementation for Artificial Intelligence applications

#### Performance boost in Image Classification

Finally, we tested whether incorporating a pre-processing WTA layer into deep learning models, such as Vision Transformers (ViTs), could enhance performance on real-world vision tasks. Specifically, we explored the role of sWTA computations in spatial feature extraction for object classification tasks. We developed a novel neural layer inspired by the hardware-constrained sWTA motif and integrated it into the conventional ViT architecture to assess its impact on classifying unseen digit datasets. This approach (Appendix A4) leveraged sWTA as a pre-processing layer to reduce redundancies and enhance contrast in visual inputs. A sliding window-based computation (**Figure 4A**) was employed for feature amplification, minimizing domain shifts (**Figure 4B**). This allowed parameters extracted by the TN hardware constraints to execute sWTA computations using recurrent excitation and lateral inhibition across pixels. In this setup, the patch with the highest variance, or ffwinner patchff, received the highest normalized value, while other patches were scaled accordingly. The selection of the ffsalientff patch size was optimized to maintain stable circuit dynamics in line with TN hardware parameters.

**Figure 4:**
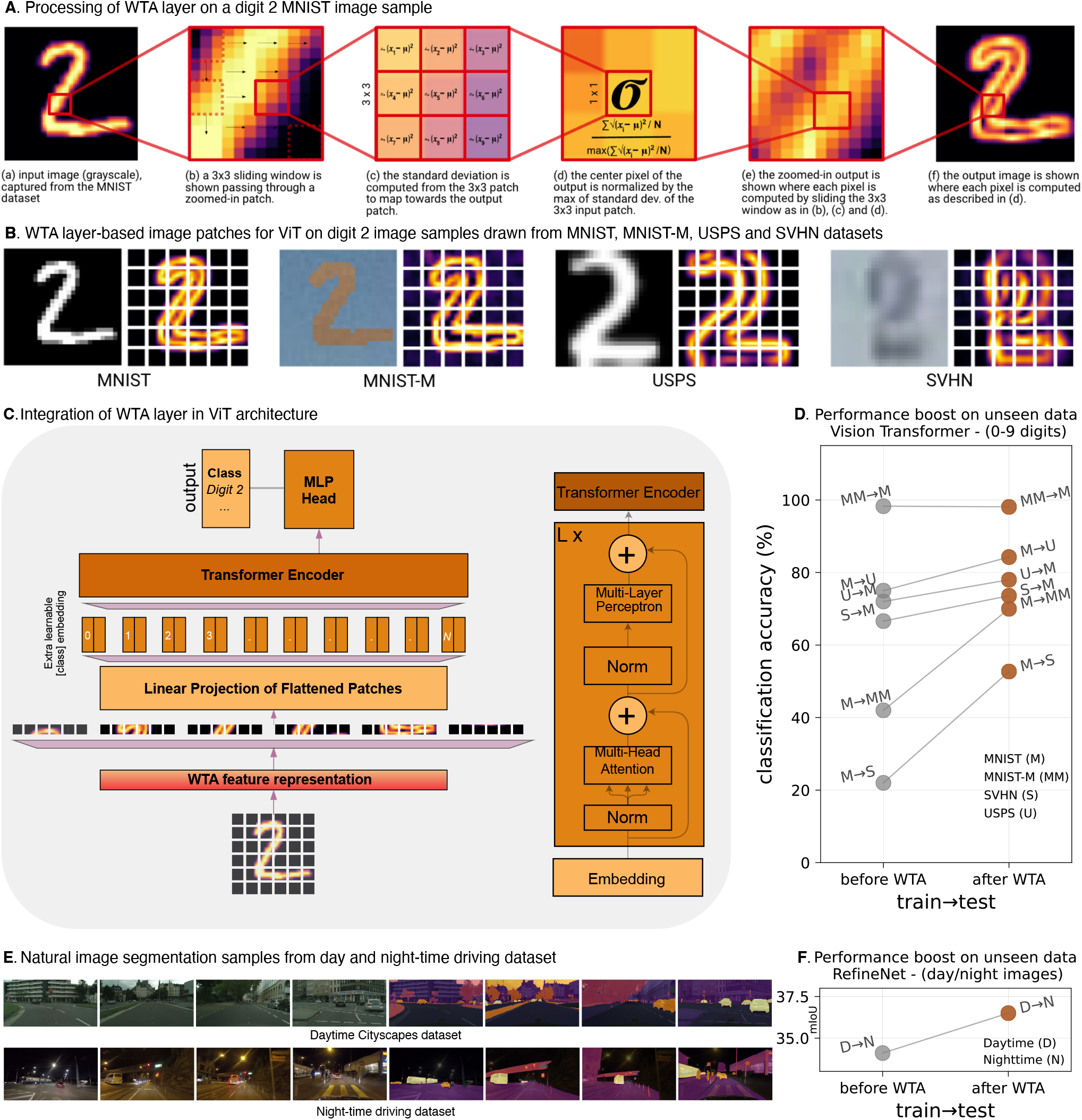
Neocortex-inspired algorithms for machine learning application and basis for zero-shot learning. A) The process involves a sliding window-based computation to enhance features, employing adjustable parameters that emulate the WTA implementation. This is accomplished through recurrent excitation and lateral inhibition acting across pixels, facilitating the feature enhancement mechanism. B) WTA layer-based image patches for ViT architecture for MNIST and digit datasets. C-D) Integration of WTA layer in Vision Transformer architecture for MNIST object classification task and results on training the model on source domain and testing on unseen target domains. E) Natural image segmentation samples from daytime and nighttime driving datasets. F) Performance improvement with and without adding a WTA layer in RefineNet for semantic segmentation task.

We next evaluated domain generalization to assess the ability of the sWTA model to adapt to unseen data distributions — an ideal test for mimicking the brain’s ability to generalize across diverse sensory inputs and maintain robust performance in novel environments. We trained the ViT model, with and without the WTA layer (**Figure 4C**), on a single source domain, and then tested its performance on unseen target domains. Significant improvements were observed across all source/target combinations (**Figure 4D**). Beyond ViT, similar performance gains were observed in EfficientNet (Tan and Le [2019]), CapsuleNet (Sabour et al. [2018]), MobileNet-V2 (Sandler et al. [2018]), and ResNet-50 (He et al. [2016]), where the WTA layer enhanced the models’ ability to learn generalizable features, improving domain shift robustness in object recognition tasks for MNIST and other digit datasets (Table 1). **Figure 4B** illustrates how the WTA layer reduces domain shift, showing high similarity across sample images post-processing. These results were achieved without intensity-based augmentation, using only geometric augmentations. EfficientNet, MobileNet, and ResNet-50 were initialized with pre-trained ImageNet weights (Russakovsky et al. [2015]). Supplementary Figures 5A-B present train/loss curve examples for ViT and CapsuleNet, while Supplementary Table 2 details model architectures and training settings. Table 1 summarizes the results, with Supplementary Table 1 showing performance improvements compared to state-of-the-art models. We also compared the WTA layer’s ability to minimize domain shift against traditional pre-processing techniques such as Local Response Normalization (LRN; Krizhevsky et al. [2017]), Local Contrast Normalization (LCN; Placidi and Polsinelli [2021]) and Z-score normalization. Supplementary Figure 6A provides qualitative comparisons, while Supplementary Figure 6B visualizes UMAP embeddings of MNIST and MNIST-M datasets post-normalization. Our WTA implementation significantly minimized domain shift (Supp Fig. 6B), leading to a marked improvement in classification accuracy (0.7) on unseen test data compared to baseline techniques (Supp Fig. 6C).

**Table 1.** Comparison of classification accuracy between WTA-based DNN architectures (Vision Transformer, EfficientNet, CapsuleNet, MobileNet, and ResNet) and source-only models trained on the MNIST (M), MNIST-M (MM), SVHN (S), and USPS (U) datasets with respective combinations as highlighted on top of each column. The top panel shows the performance of the models without adding the WTA layer whereas the bottom panel shows the performance boost by adding the WTA layer to the network architectures. The models are tested on completely ‘unseen’ target datasets. (**bold-red** indicates the best and **bold-black** indicates the 2nd best)

| Source-only Models | M→U | U→M | S→M | M→S | M→MM | MM→M |
| --- | --- | --- | --- | --- | --- | --- |
| ViT | 75.0 | 72.0 | 66.6 | 22.0 | 42.0 | 98.3 |
| EfficientNet | 77.9 | 50.7 | 61.4 | 18.6 | 18.8 | 95.0 |
| CapsuleNet | **96.4** | **87.2** | 58.1 | 11.8 | 22.5 | **98.4** |
| MobileNet | 84.4 | 60.0 | 72.2 | 22.4 | 33.9 | 97.3 |
| ResNet | 82.5 | 58.5 | 63.4 | 27.2 | 38.2 | 97.4 |
| ViT+WTA | 84.26 | 78.0 | **73.6** | **52.7** | 70.0 | 98.1 |
| EfficientNet+WTA | 83.5 | 74.2 | 69.1 | 19.6 | 48.8 | 96.3 |
| CapsuleNet+WTA | **94.1** | **87.8** | **75.9** | 32.1 | 57.2 | **98.6** |
| MobileNet+WTA | 82.5 | 70.9 | 73.5 | **40.5** | **73.4** | 97.8 |
| ResNet+WTA | 82.8 | 66.0 | 71.2 | 27.7 | **70.2** | 97.8 |

#### Performance boost in Image Segmentation

Lastly, we evaluated our approach in a deep learning model for the challenging task of natural image segmentation. Similar to the results seen in image classification, incorporating the sWTA layer into the RefineNet architecture (Lin et al. [2017]) with ResNet-101 significantly improved performance in semantic segmentation. This underscores the broad applicability of our approach across various vision tasks. The model was trained on the Cityscapes dataset (Cordts et al. [2016]), consisting of 2,975 daytime driving images, and tested on 50 coarsely annotated Nighttime Driving dataset (Dai and Van Gool [2018]) (**Figure 4E**). We employed the same training setup, using a dynamic learning rate of 0.1 with stochastic gradient descent (SGD) on an RTX A6000 GPU and a batch size of 6. Performance was measured using mean Intersection over Union (mIoU).

Notably, adding the WTA layer to RefineNet achieved performance on par with nighttime driving data using only source-trained models (**Figure 4F**).

Table 2 highlights the performance improvements in RefineNet for natural image segmentation tasks, with and without the sWTA layer, demonstrating its robustness in handling domain shifts. Supplementary Fig. 7 shows qualitative results for the nighttime dataset, comparing performance with and without the WTA layer.

**Table 2.** Segmentation performance on Nighttime Driving (Dai and Van Gool [2018]), reported as mIoU scores. WTA-RefineNet outperforms other methods trained only on daytime data and has a competitive performance to methods also using nighttime images

| Method | Nighttime Driving (mIoU) |
| --- | --- |
| RefineNet Lin et al. [2017] | 34.1 |
| W-RefineNet Lin et al. [2017]<br>(pre-trained RGB weights of Resnet on ImageNet) | 34.6 |
| RefineNet-AdaBN Lin et al. [2017] | 36.3 |
| WTA-RefineNet<br>(pre-trained RGB weights of Resnet on ImageNet) | **36.5** |

In conclusion, implementing sWTA motifs as a layer in various deep learning architectures substantially improves their performance. Using the competitive-cooperative dynamics of biologically inspired WTA mechanisms as a preprocessing layer creates a synergy between biological principles and artificial neural networks. Our WTA-inspired layer enhances performance in real-world tasks like image classification and segmentation by acting as an adaptive filter that sharpens focus on the most relevant features while reducing noise and irrelevant information.

## Discussion

The primary objective of this study was to translate core neurobiological principles into computational frameworks that can be efficiently implemented on neuromorphic hardware and modern AI/ML models such as transformers. Specifically, we focused on the soft winner-take-all (sWTA) mechanism, a fundamental computational motif observed in neural circuits. Rather than implementing detailed, complex neurobiological circuits with a one-to-one mapping of circuit elements onto hardware, our goal was to extract the core computational principles based on a biophysically realistic cortical circuit architecture that captures common principles across sensory cortical regions and translate them efficiently into neuromorphic and AI models.

To validate this approach, we first examined experimentally constrained biophysical models to confirm that biologically realistic cortical circuits indeed implement sWTA computations. We designed a canonical cortical circuit consisting of distinct interneuron subpopulations, where the proportions of these neurons were selected based on prior experimental observations (Pfeffer et al. [2013b]). Unlike models that rely on non-biophysically realistic spiking neuron approximations (Potjans and Diesmann [2014]), our model incorporated Hodgkin-Huxley-based conductance mechanisms, ensuring biological plausibility. The synaptic weights in the biophysical circuit model were obtained from our experimental data. This enabled us to develop a mechanistic model that captures the core cortical circuit primitives, validates computational principles such as sWTA, and extracts the contribution of individual circuit components in terms of distinct interneuron types. These validated circuit elements were then systematically implemented in neuromorphic hardware and AI/ML models.

This foundational step ensured that our framework was grounded in biologically plausible mechanisms before adapting it for hardware implementation. We then explored how WTA computational primitives could be leveraged to construct neural state machines on IBM’s TrueNorth (TN) neuromorphic hardware. Finally, we investigated the integration of hardware-derived parametric constraints and computational principles extracted from the biophysical model into deep learning models, such as Vision Transformers, to assess their impact on training efficiency. Our gain-matching technique facilitates the emulation of diverse neural network architectures and computational principles on neuromorphic hardware, building on prior research in analog systems (Neftci et al. [2011]). This work extends the application of sWTA networks, which have been explored in contexts such as object recognition (Yuille and Grzywacz [1988], Riesenhuber and Poggio [1999], Erlhagen and Schöner [2002]), attention (Itti et al. [1998], Deco and Rolls [2005]), and decision-making (Amari [1977]). Additionally, our framework could be adapted for next-generation neuromorphic platforms like Braindrop Neckar et al. [2018b], IBM’s Northpole (Modha et al. [2023b]), and Intel’s Loihi2 (Davies et al. [2021]). However, further empirical validation is necessary to confirm the extent to which these insights can be generalized across different neuromorphic architectures.

Building on prior studies, we explored how generic circuit motifs used for sensory computations could be adapted for working memory in cognitive decision-making tasks on TN neuromorphic hardware (Neftci et al. [2013b], Rutishauser and Douglas [2009b]). We found that our approach allows for efficient scaling of state machines. However, further comparative analysis with alternative architectures is required to comprehensively evaluate its performance. Our work offers a cortical computation primitive-based approach where complex algorithms could be implemented by building on simplified computational primitives. Previous work (Pei et al. [2019]) has demonstrated that convolutional and spiking neural networks (SNNs) have also been efficient for neuromorphic hardware and constructing artificial cognitive agents. However, the architecture used in this study differs from those found in the mammalian brain. Future work could explore integrating our biologically inspired computational primitives with large-scale SNNs to develop artificial general intelligence. Additionally, investigating the role of specific interneurons and circuit motifs beyond sensory regions could reveal additional principles relevant to context-dependent decision-making. Extending our framework to incorporate dendritic computations and related learning rules would require further modeling and experimental validation.

In summary, our study highlights how fundamental neurobiological principles, particularly sWTA, can be translated into scalable neuromorphic and AI/ML implementations (Supplementary Figure 1). By effectively capturing the computational principles and efficiency of core neurobiological components, we provide a framework that bridges theoretical neuroscience, neuromorphic hardware, and AI. Future refinements and empirical studies could further enhance this approach, contributing to the broader advancement of NeuroAI.

## Acknowledgments

We thank Ehsan Arabzadeh, William Connelly, and Giacomo Indiveri for the fruitful discussion. We thank Jan Drugowitsch, Christopher Harvey, Debanjan Dasgupta, Alessandro Galloni, and Samuel Gershman for their helpful comments and constructive criticism of the manuscript. We thank Dharmendra Modha, Hayley Hu, and Ben Shaw for assistance and guidance on the TrueNorth hardware. The neuromorphic implementation was performed at the Institute of Neuroinformatics (INI), UZH/ETH Zurich as a part of the IBM-INI collaboration. We thank the organizers of the Telluride workshop and IBM Bootcamp. We acknowledge the generous support of high-computing GPUs from Tibbling Technologies for running experiments to train and test the WTA implementation in deep learning architectures.

## Author Contributions

A.I., G.F., and S.H. designed research and wrote the manuscript; S.H. performed *in-vitro* experiments, biophysical modeling, and neuromorphic hardware implementation; A.I. and S.H. conceived mapping of neocortical computation in deep learning architectures. A.I. and H.M. performed mapping of WTA in deep learning architectures and ran experiments for image classification and segmentation tasks; G.J.S supervised *in-vitro* experiments; G.F supervised biophysical modeling and contributed to the organization of the manuscript.

## Methods

### Animals

All experimental procedures were approved by and conducted in accordance with Harvard Medical School and The Australian National University Institutional Animal Care and Ethics Committee.

### Viral injections and Whole-cell patchclamp recordings

For labeling bottom-up sensory input and top-down contextual inputs, AAV1-hSyn-hChR2(H134R)-EYFP-WPRE-hGH (ChR2) was injected in either the contralateral visual cortex (or visual thalamus, dLGN) and primary auditory cortex respectively (Honnuraiah et al. [2024]; Godenzini et al. [2021]). Ipsilateral eye input is stimulated by contralateral V1, and contralateral eye input is stimulated by dLGN stimulation ([Honnuraiah et al. [2024]). For PV and SST inhibition experiments, Cre-dependent ChR2 (AAV1-EF1a-DIO-hChR2(H134R)-EYFP-WPRE-hGH) was injected in the binocular visual cortex of the transgenic mice expressing Cre in either PV or SST. Three to four weeks after viral injection mice were deeply anesthetized with isoflurane (3% in oxygen) and immediately decapitated. Slice preparation protocols and the experimental recordings are explained in detail in this study (Honnuraiah et al. [2024]). All recordings were made in the current-clamp using a current clamp BVC-700A amplifier (Dagan Instruments, USA). Data was filtered at 10 kHz and acquired at 50 kHz by a Macintosh computer running Axograph X acquisition software (Axograph Scientific, Sydney, Australia) using an ITC-18 interface (Instrutech/HEKA, Germany). Hyperpolarizing and depolarizing current steps (200 pA to +600 pA; intervals of 50 pA) were applied via the somatic recording pipette to characterize the passive and active properties of neurons. Brain slices were bathed in gabazine (10 µM) to block inhibition mediated by GABA-A receptors. Other pharmacological agents used in these experiments included tetrodotoxin (TTX; 1 µM) and 4-aminopyridine (4-AP; 100 µM), as noted in the Results. For photo-stimulation of ChR2-expressing neurons and axon terminals, a 470 nm LED (Thorlabs) was mounted on the epi-fluorescent port of the microscope (Olympus BX50) allowing wide-field illumination through the microscope objective. The timing, duration, and strength of LED illumination were controlled by the data acquisition software (Axograph).

### Computational Modeling

We conducted biophysical modeling using the NEURON 8.2 simulation environment (Carnevale and Hines [2006], Hines and Carnevale [1997]), with an integration time constant of 25 µs. The active and passive properties of the model were optimized to match the experimental recordings (Supp Fig 1). We set the passive parameters as follows: Internal/axial resistance (Ri/Ra) to 150 #.cm, membrane resistance (Rm) to 30 K#.cm2, capacitance (Cm) to 1 µF/Cm2 and resting membrane potential (Vm) to -75 mV. All neurons were simplified and implemented as a “ball and stickff model consisting of a somatic compartment (dimensions: Length=50 µm; diameter=50 µm) and a single dendritic compartment (dimensions: Length=100 µm; diameter=1 µm). Dendritic compartments were passive and were not adjusted for spines in the interneuron population but were adjusted for spines in pyramidal neurons by scaling the Cm by 2 and Rm by 0.5. Active conductances were included in the somatic compartment to mimic the regular firing pattern of pyramidal neurons, fast-spiking pattern of PV, burst-spiking for SST, and delayed spiking from the Lamp5. Active ion-channel distribution and its conductance values are obtained from our previous study (Soldado-Magraner et al. [2020]). A synapse was modeled as a co-localized combination of NMDA and AMPA receptor currents. A default value of NMDAR:AMPAR ratio was set at 1.5. All the values related to synaptic parameters were obtained from our previous study (Narayanan [2013],

### Rate-based abstract neural network model

We developed a simplified abstract model to reduce computational demands and extract the principles from detailed biophysical network models. We have used a rate-based approach to model neuronal activity. We approximate the neuronal activation by a linear-threshold function that describes the output action potential discharge rate of the neuron as a function of its input Bauer et al. [2014]. This type of neuronal activation function is a good approximation to experimental and biophysical observations of the frequency of action potential discharge to synaptic or current inputs. The change in activity of a neuron is modeled as the summation of synaptic input with a decay of the current activity. The dynamics of the activity of the rate-based neurons implemented are given below:

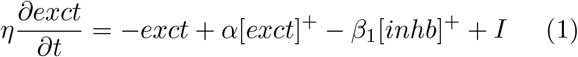

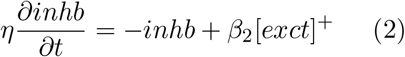

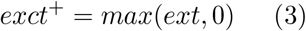

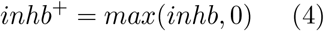

Where, [exc_*t*_]^+^ is excitatory neuron activity, [inh_*t*_]^+^ is inhibitory neuron activity, and τ is the neuronal time constant, α, β_1_, and β_2_ are synaptic weights (Figure 2B).

The implementation of the computational primitives obtained from the biophysical models in the rate-based abstract models was crucial. This is because it provided analytically tractable solutions for the dynamics of neural activity. The analytical solution was later used to derive constraints for the TN hardware parameters as explained in Appendix A3.

### IBM TrueNorth hardware and Compass software emulator

The IBM TN neuromorphic chip is composed of 64×64 [4096] digital neurosynaptic cores tiled in a 2-D array, containing an aggregate of 1 million neurons and 256 million synapses. Each core implements 256 neurons single-compartment leaky-integrate-and-fire neurons which could be operated in either rate or spike mode configuration. Each core is supported by a 256×256 crossbar synapse array, and communication circuits to transfer spike trains. The crossbar array is flexible and can be configured freely. Each row of the crossbar corresponds to an axon of the neuron represented by horizontal lines which could be driven by any on-chip neurons. Inputs to the cores are generated by configuring the on-chip neurons to generate various frequencies either in rate or spike mode. Each column corresponds to a dendrite of that particular neuron represented by a horizontal line. A connection between an axon and a dendrite is a synapse and is organized into a synaptic crossbar (Supplementary Figure 4A). A peripheral memory core is located at the intersection of each row and a column, and the binary value stored in the core represents whether or not a connection exists between the particular axon-dendrite pair. Therefore, each neuron can be configured to receive up to 1024 synaptic inputs (through its dendrite) depending on the crossbar value and the activity of the axons. TN operates in a mixed asynchronous–synchronous approach. All the communication and control circuits operate in asynchronous design while computations are done in synchronous design. Since TN cores operate in parallel and are governed by spike events, it is natural to implement all the routing mechanisms asynchronously. All the core computations must finish with finish in the current tick which spans 1 ms. Compass is the software emulator to program and simulate the full 4096 neurosynaptic cores and the digital asynchronous–synchronous design ensures one-to-one compass to TN correspondence (Akopyan et al. [2015] Merolla et al. [2014]).

### Vision Transformer (ViT) architecture

Inspired by the original transformer ([Vaswani [2017]) architecture for Natural Language Processing, ViT (Dosovitskiy et al. [2020]) is a self-attention-based architecture. It works as follows: the input image is distributed into N (flattened 2D) patches (where we keep N=6 for the MNIST experiments) and linear embeddings of these patches are fed as an input to the encoder of the trained ViT. The image patches of digits are embedded as tokens. The encoder block has several multi-headed layers with self-attention along with a normalization layer at the start of each layer. Furthermore, a Multi-Layer Perceptron (MLP) with a single hidden layer is used as a classification head that predicts the object categories present in the input image.

### WTA implementation in ViT

We present a Winner-Take-All (WTA) approximation as a neuro-inspired layer in the Vision Transformer (ViT) architecture. Inspired by distinctive properties of cortical circuits in the mammalian visual cortex, this layer captures the neural signal regulation characteristics. A defining feature of these neurons lies in their ability to intricately capture and encode the contrasting elements and structural nuances of visual stimuli. This capability is reflected in the variable neural firing sequence of the neurons that WTA emulates, aligning closely with the varying contrasts present in the stimuli. This results in a more refined and contextually aware representation, which is particularly beneficial in object classification contexts where adaptability and nuanced understanding of visual stimuli are crucial. Mathematical implementation is described in Appendix A4.

### MobileNet with WTA

Developed for embedded devices such as mobile phones, etc., MobileNet-v2 (Sandler et al. [2018]) is successfully reducing the number of parameters by depth-wise separable convolutions, while keeping the accuracy comparable to the state-of-the-art. We initialized the weights of MobileNet-V2, trained on ImageNet, and added the WTA layer to the model.

### EfficientNet with WTA

EfficientNet-B0 (Tan and Le [2019]) is used with pre-trained weights for ImageNet. It is pre-trained to classify 1000 image classes and trained on more than a million images. We initialized the convolution layers with the pre-trained model weights. We added the WTA layer after the input layer into the model.

### ResNet with WTA

Among different variants of ResNet, we select ResNet-50 (He et al. [2016]), which contains 50 neural network layers. The introduction of skip connections reduces the problem of vanishing gradient and also ensures that the higher layers do not perform any worse than the layers before by learning the identity function. Similar to the other models above, we are initializing the weights with the pre-trained model on ImageNet and including the WTA layer after the input layer in the model architecture.

### CapsuleNet with WTA

Unlike other Convolutional Neural Network (CNN) architectures, CapsuleNet (Sabour et al. [2018]) applies pattern matching by decomposing the hierarchical representations of the input features. The eventual representation of this network is supposed to be invariant to the view-angle of the input samples. One of the major differences between a typical CNN and CapsuleNet is the output of the individual units in their architecture. While the output of a single neuron in a CNN is mostly a scalar value, it is a vector in the case of CapsuleNet. Similar to the ViT, we include the WTA layer as an initial layer in the CapsuleNet architecture to make it robust for domain adaptation tasks for digit datasets.

### RefineNet with WTA

RefineNet (Lin et al. [2017]) is a versatile multi-path refinement network that leverages all the information gathered during the down-sampling process to facilitate high-resolution prediction through long-range residual connections. This approach enables the deeper layers, which capture high-level semantic features, to be directly refined using fine-grained features from earlier convolutions. The individual components of RefineNet employ residual connections following the identity mapping principle, enabling efficient end-to-end training. In our experiments, we used pre-trained RGB weights of ResNet on ImageNet for training and testing RefineNet with and without adding a WTA layer.

### MNIST and digit datasets

For the domain generalization task, we utilized a suite of digit datasets that included the MNIST, SVHN, USPS, and MNIST-M (USPS, and MNIST-M (LeCun et al. [2010], Netzer et al. [2011], Hull [1994], Ganin et al. [2016]). Each dataset was split into 70-20-10 train, val, and test

**MNIST dataset**, introduced by LeCun et al. [2010], is one of the most widely used datasets for handwritten digit classification. It contains a total of 70,000 grayscale images of handwritten digits. Each image is of size 28×28 pixels, and the dataset has been instrumental in benchmarking various machine learning algorithms.

**SVHN (Street View House Numbers) dataset**, presented by Netzer et al. [2011] is a real-world image dataset obtained from house numbers in Google Street View images. It comprises over 600,000 digit images. Specifically, it contains 73,257 digits for training, 26,032 digits for testing, and an additional 531,131 somewhat less difficult samples that can be used as extra training data. This dataset challenges models with recognizing digits in more complex and varied scenarios compared to the controlled environment of MNIST.

**USPS (United States Postal Service) dataset**, introduced by Hull [1994] is another hand-written digit dataset used for text recognition research. It contains 9,298 16×16 grayscale images of handwritten digits. The dataset was derived from scanned mail and has been a staple in the handwritten digit recognition field.

**MNIST-M dataset**, presented in the work by Ganin et al. [2016], is a modified version of the original MNIST dataset. It was created by overlaying MNIST digits onto patches randomly extracted from color photos of the BSDS500 dataset (Arbelaez et al. [2010]), resulting in a blend of digits and colored backgrounds. The MNIST-M dataset contains 149,002 images. This combination introduces additional challenges due to the color and texture variations in the background, making it a valuable dataset for studying domain adaptation.

For domain generalization tasks, these datasets are particularly valuable because they offer variations in terms of image quality, resolution, and real-world applicability. The diversity in these datasets, ranging from clean handwritten digits to digits in natural scenes, challenges models to generalize well across different domains. This makes them ideal benchmarks for evaluating the robustness and adaptability of machine learning algorithms, especially in scenarios where the training and test data distributions differ significantly.

### Natural image dataset

To explore the effect of WTA for segmentation tasks on natural images, we select a cityscape Cordts et al. [2016] data for training and nighttime driving Dai and Van Gool [2018] dataset for testing. This evaluation aims to test the robustness of the model against day-to-night time domain shifts.

## Appendix

### 0.1 Emulating Cortical Neuron Physiology in TN Hardware

#### Dynamic Range Relationship

The dynamic range *r* of the neuron is given by:

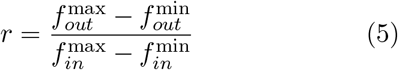

This equation expresses how the output firing rate scales with the input firing rate.

#### Sensitivity of a Single Synaptic Weight

We define the sensitivity of a single synaptic weight:

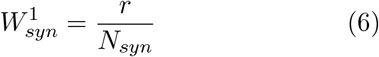

where:

- *N*_*syn*_ is the total number of synapses,
- 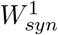represents the weight sensitivity per synapse.

This ensures that the effect of a single synapse is properly scaled based on the number of synapses contributing to the neuron’s response.

#### Maximum and Minimum Crossbar Synaptic Weight

The range of the crossbar synaptic weight is given by:

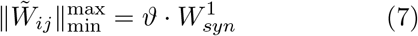

where:

- ϑ is a scaling factor,
- 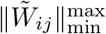represents the spread of crossbar synaptic weight values.

This ensures that synaptic weights remain within a controlled range.

#### Leak Parameter of TN Neuron

The leak parameter λ is determined by:

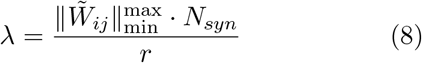

This establishes a relationship between the synaptic weight range, total synapses, and dynamic range, ensuring that the leak parameter is appropriately set to balance input contributions.

#### Example Calculation

Given:

- *N*_*syn*_ = 10
- *r* = 1
- 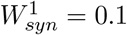
- 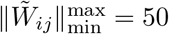

We compute:

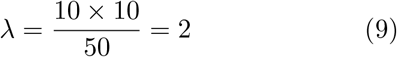

This threshold setting ensures that the neuron’s behavior remains within physiological ranges.

#### Excitatory and Inhibitory Synaptic In-Teractions

To incorporate excitatory and inhibitory synaptic contributions:

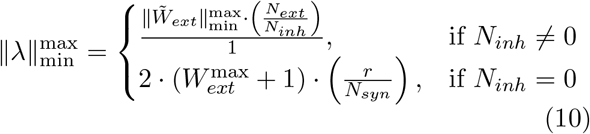

This ensures a balance between excitatory and inhibitory influences in synaptic integration.

Thus, by carefully tuning parameters ϑ, λ, and 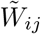, we achieve biologically realistic synaptic integration, closely resembling cortical neuron behavior.

### 0.2 Hardware load analysis

To analyze the computational load for implementing the neuromorphic sWTA model, we define the following computational parameters:

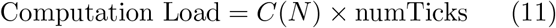

- *C*(*N*) represents the number of connector pins required,
- numTicks denotes the total computational time steps.

The total computational load *C*(*N*) can be decomposed into input and output loads:

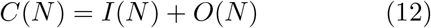

where:

- *I*(*N*) is the number of input pins,
- *O*(*N*) is the number of output pins.

By optimizing the architecture of the neuromorphic hardware, we aim to minimize the number of connector pins while ensuring efficient signal processing and minimal latency in winner-take-all (sWTA) operations.

### 0.3 Automated Parameter Mapping to IBM TN Parameters and Derivation of Stability Constraints

In this section, we provide derivation of the TN neuron model parameters, incorporating leak, threshold, excitatory, and inhibitory feedback. We then derive stability constraints via contraction analysis to implement winner-take-all behavior on IBM TN hardware. TN neurons are configured to operate as linear threshold units (in rate-based mode) as described in the previous section.

Based on this linear operation, we can estimate the role of the self-excitatory feedback connection on the transfer function, according to the equation below: let’s start with the simple case without leak and threshold and progressively add these variables.

Assuming no leak (λ = 0) and no threshold (θ = 0), the output firing rate is:

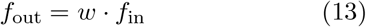

This represents a pure linear relationship, where the neuron’s response is an amplified version of the input. To account for decay in neural activity over time, we introduce the leak parameter (λ):

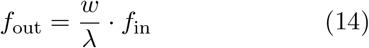

where λ acts as a damping factor, reducing the effective influence of the input.

Neurons typically require a minimum level of activation before they fire, which is represented by the threshold (θ). A minimum input activation threshold (θ) modifies the transfer function to:

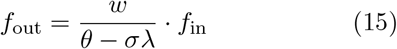

ensuring that unless the input reaches a certain level, the neuron remains inactive.

Based on our TN parameter sensitivity analysis **Supplementary Figure 3A-C**, we conclude that the threshold (θ) and leak (λ) parameters in TN reflect PV and LAMP5 interneuron mediated inhibition in our biophysical model respectively. While the common inhibitory population captures the SST-mediated inhibition.

To model realistic neural circuits, we incorporate feedback connections that are mediated by distinct excitatory and inhibitory pathways:

- **Excitatory recurrent feedback** (α) amplifies the neuron’s response.
- **Inhibitory feedback** (β_1_, β_2_) suppresses the neuron’s response.

The updated transfer function incorporating feedback mechanisms is:

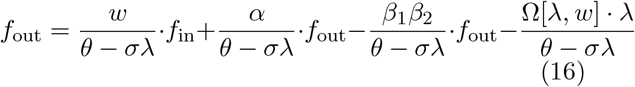

Rearranging:

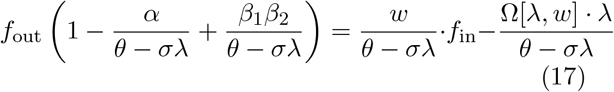

Simplifying further:

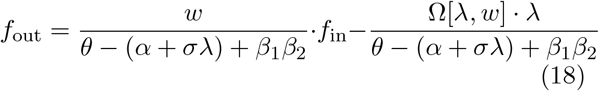

Defining the gain function:

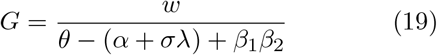

This represents the effective scaling factor for the neuron’s response, incorporating both excitatory and inhibitory effects.

For global stability, we use contraction analysis, which requires the system to satisfy:

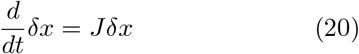

where stability is ensured if:

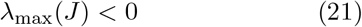

Computing the Jacobian:

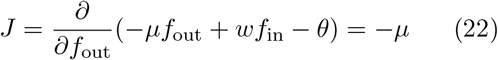

For contraction stability:

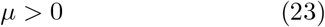

With feedback:

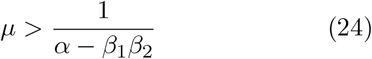

This constraint ensures that the system does not become unstable due to excessive excitatory feedback and maintains the system gain (*G*) in stable bounds.

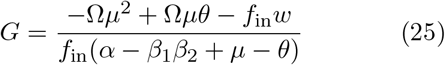

where Leak-Weight Interaction Function:

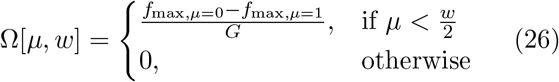

For winner-take-all behavior to emerge, additional constraints are required:

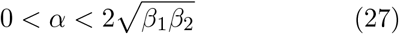

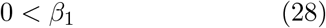

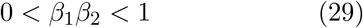

These constraints ensure that only the most strongly activated neuron dominates, suppressing the responses of weaker neurons.

Optimal parameters satisfying these constraints:

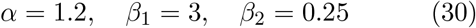

with corresponding TrueNorth/Compass parameters:

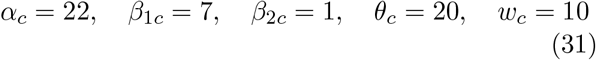

We then plug in the above values in the TrueNorth/Compass circuit shown in **Figure 3** and verify if the following WTA functional characteristics are satisfied:

- Non-linear signal amplification (winner selection), validated in **Figure 3G**.
- Robustness and signal restoration with broadly tuned inputs, validated in **Figure 3H**.
- Dynamic switching and multi-stability, validated in **Figure 3I**.

Thus, by systematically incorporating leak, threshold, and feedback, obtained from the above equations, we configure TN neurons to exhibit stable, competitive, and winner-take-all behavior that closely matches the cortical neurons.

### 0.4 Mathematical formulation of WTA in ViT

To mathematically implement the WTA layer, we process an input image *I* ∈ R^*W* ×*H*×*C*^, where:

- *W, H, C* represent the width, height, and number of channels,
- The image is divided into patches *P* = {*P*_*s*1_, *P*_*s*2_, …, *P*_*sn*_} of size *s*, each centered around pixel *k* at coordinates (*i, j*),
- The goal is to derive a biologically realistic **Winner-Take-All (WTA) computation** that emphasizes the most relevant features while suppressing irrelevant inputs.

#### Putative mean computation via SST Interneurons

The local mean over each patch is given by:

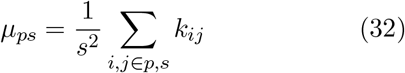

This expression represents the average pixel intensity within each patch. However, in a biologically inspired framework, the mean computation is influenced by feedback inhibition mediated by SST (somatostatin) interneurons. This modifies the expression to which in terms of **SST interneuron-based computation** translates to:

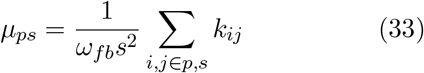

where ω_*fb*_ represents **feedback inhibition** by SST interneurons, responsible for computing local averages.

#### Putative variance computation via PV Interneurons

The variance within a patch is a crucial measure for contrast enhancement and feature detection. It is computed as:

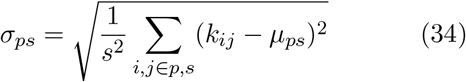

Biological circuits incorporate feedforward inhibition through PV (parvalbumin) interneurons, which adjust the gain of excitatory input. Thus, the expression transforms into:

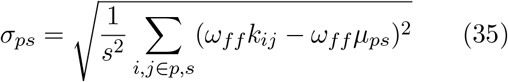

where ω_*ff*_ represents **feedforward inhibition** by PV interneurons, which selectively amplifies significant inputs.

#### Putative normalization via LAMP5 Interneurons

Normalization ensures that the strongest response is emphasized across all patches. The strongest response is selected across all patches:

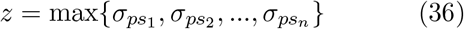

This maximum standard deviation is then used to normalize the contrast values:

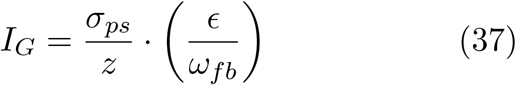

where ϵ is a small stabilization constant, and LAMP5 interneurons contribute to global gain control by regulating recurrent excitation and global gain control, ensuring the stability of feature selection.

#### Derivation inspired from biophysical models

Based on our sensitivity analysis of the biophysical model **(Supplementary Figure 2H-I)**, we derived putative computations of specific interneurons and applied them in our prefilter. Specifically, we hypothesized that:

- SST interneurons regulate averaging via feed-back inhibition.
- PV interneurons compute local contrast via feed-forward inhibition.
- LAMP5 interneurons perform global normalization via gain control.

Incorporating the phenomenologically derived biophysical constraints, we derive the final equations:

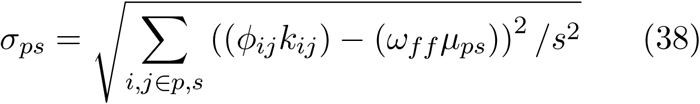

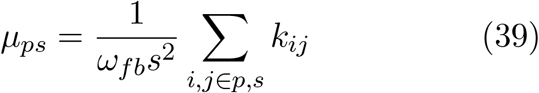

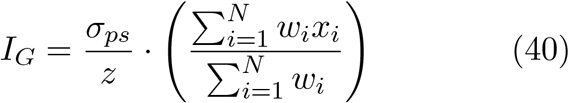

where *W*_*i*_ are synaptic weights modulating LAMP5 inhibition.

Finally, we demonstrate that **WTA computation in ViT aligns with cortical circuits computational primitive**, ensuring biologically inspired feature selection and domain adaptation.

##### Algorithm 1

Mapping Biophysical Model Parameters to Experimental Data

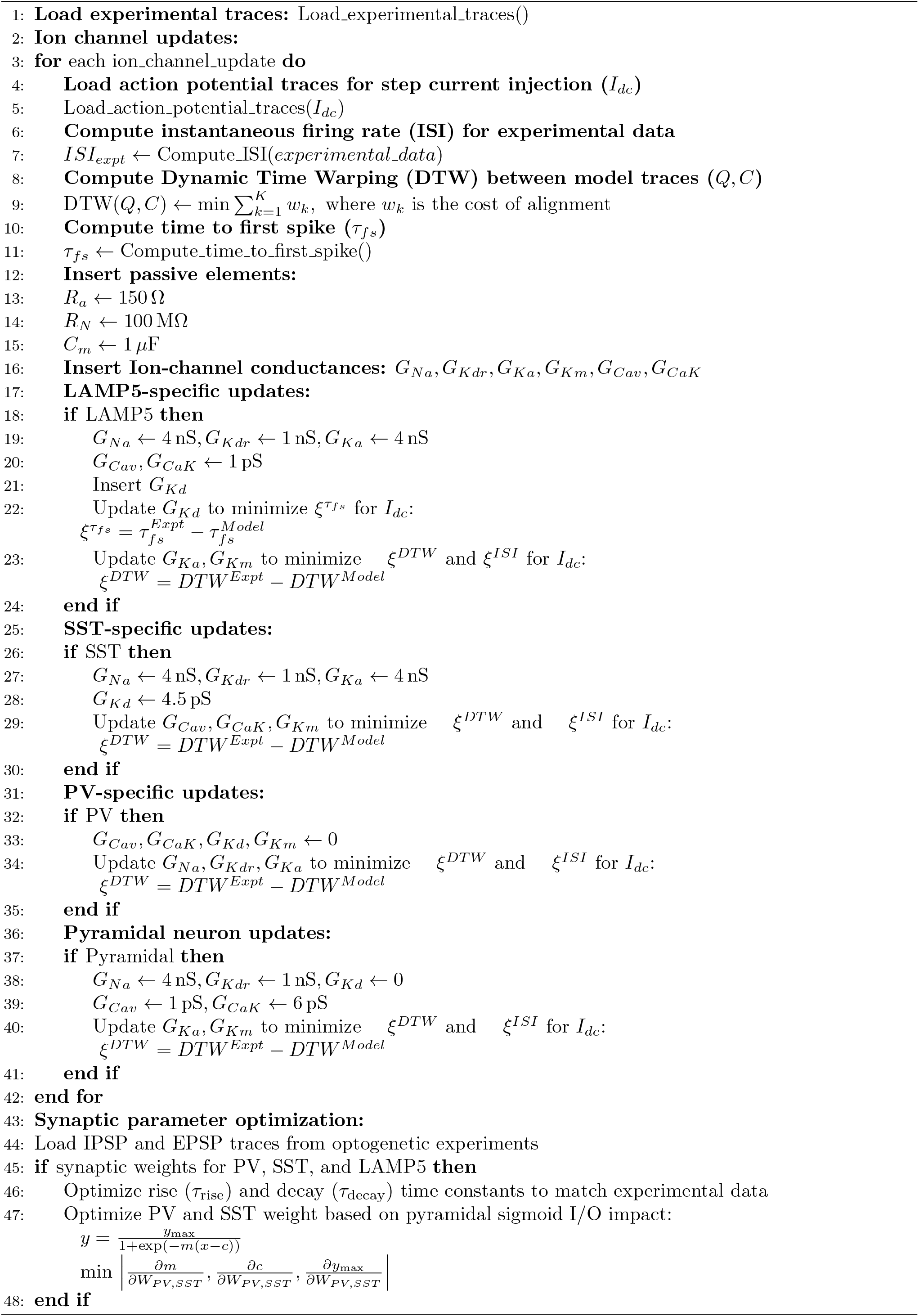

##### Algorithm 2

Mapping Biological Parameters to IBM TrueNorth Parameters

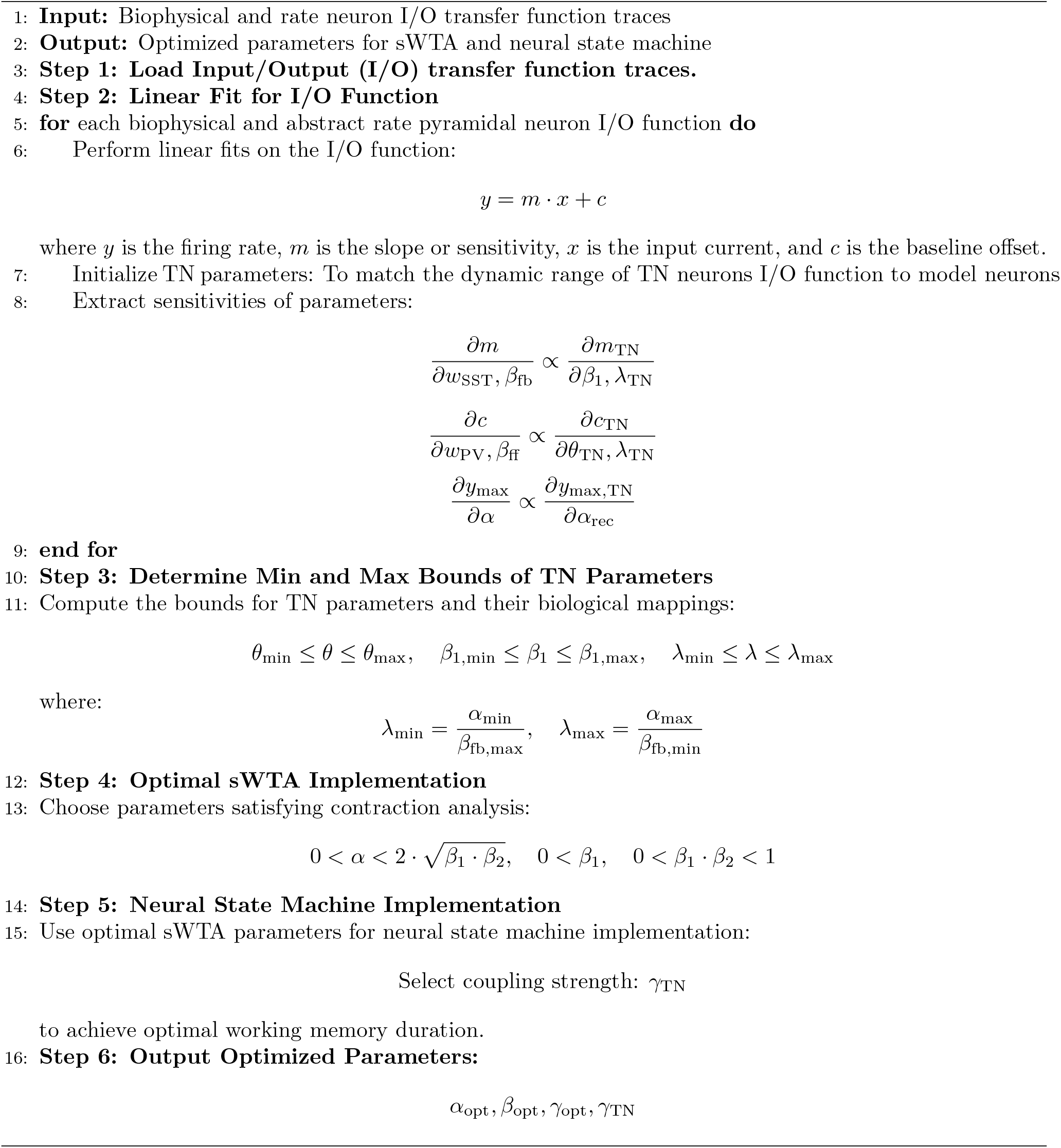

##### Algorithm 3

Mapping AI Parameters to Biological Counterparts

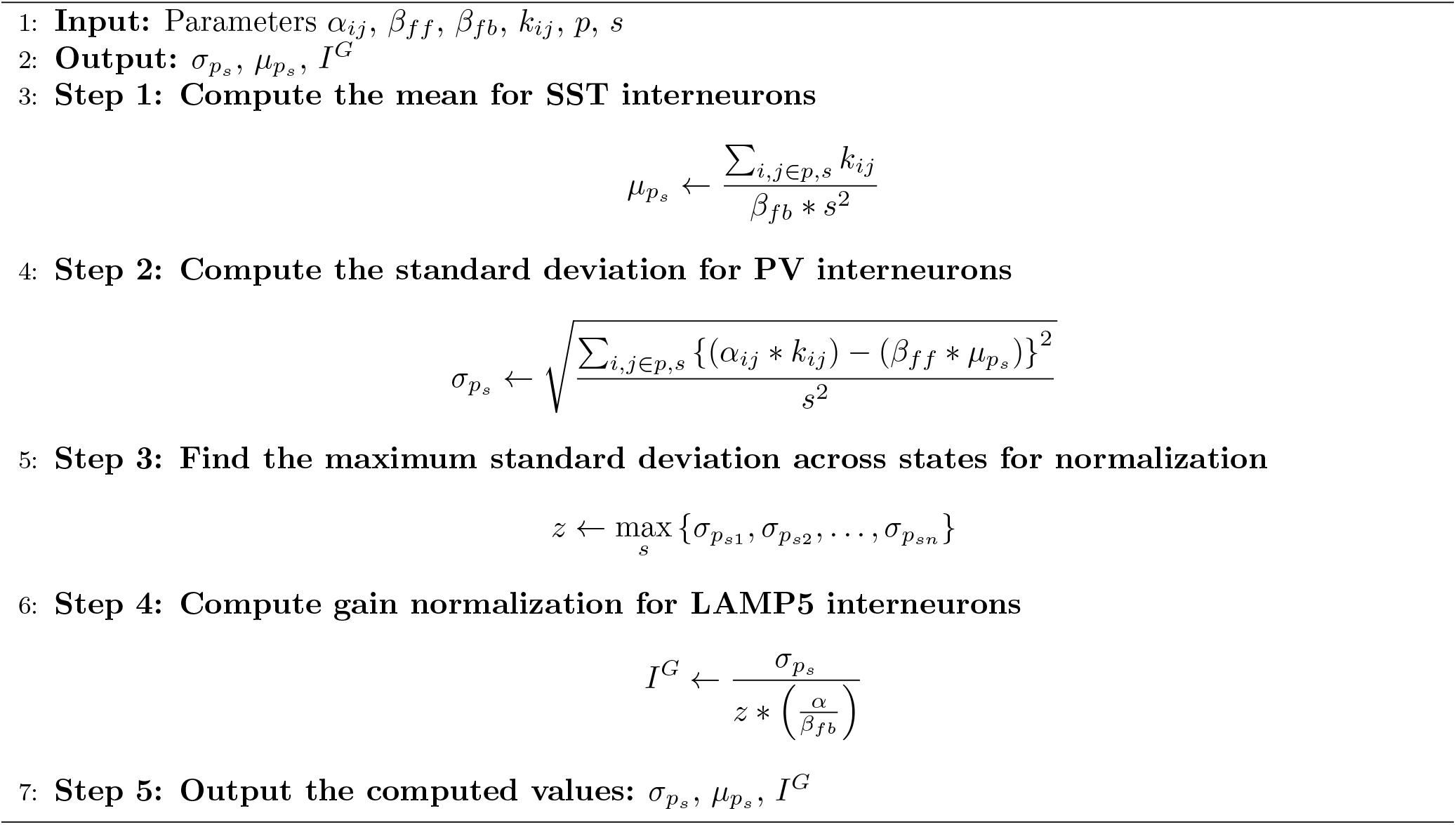

## Neuromorphic Hardware

**Supplementary Figure 1:**
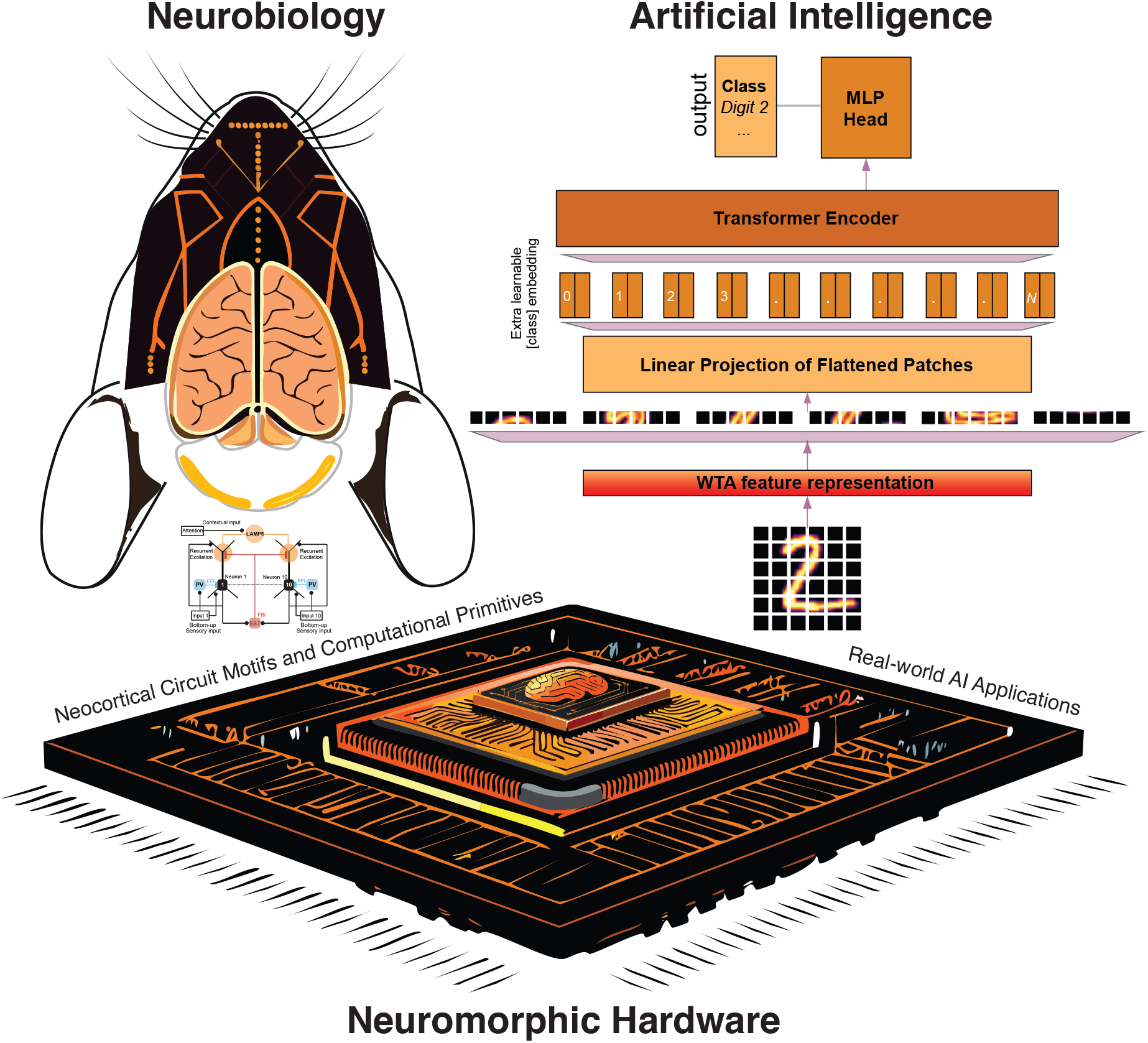
Block diagram architecture of our proposed framework. We draw inspiration from understanding the neurobiological systems of the brain and implement our findings into the neuromorphic hardware. In addition, we implement our findings as a pre-processing WTA layer in AI models and observe high-performance boosts in real-world object classification and natural image segmentation tasks.

**Supplementary Figure 2:**
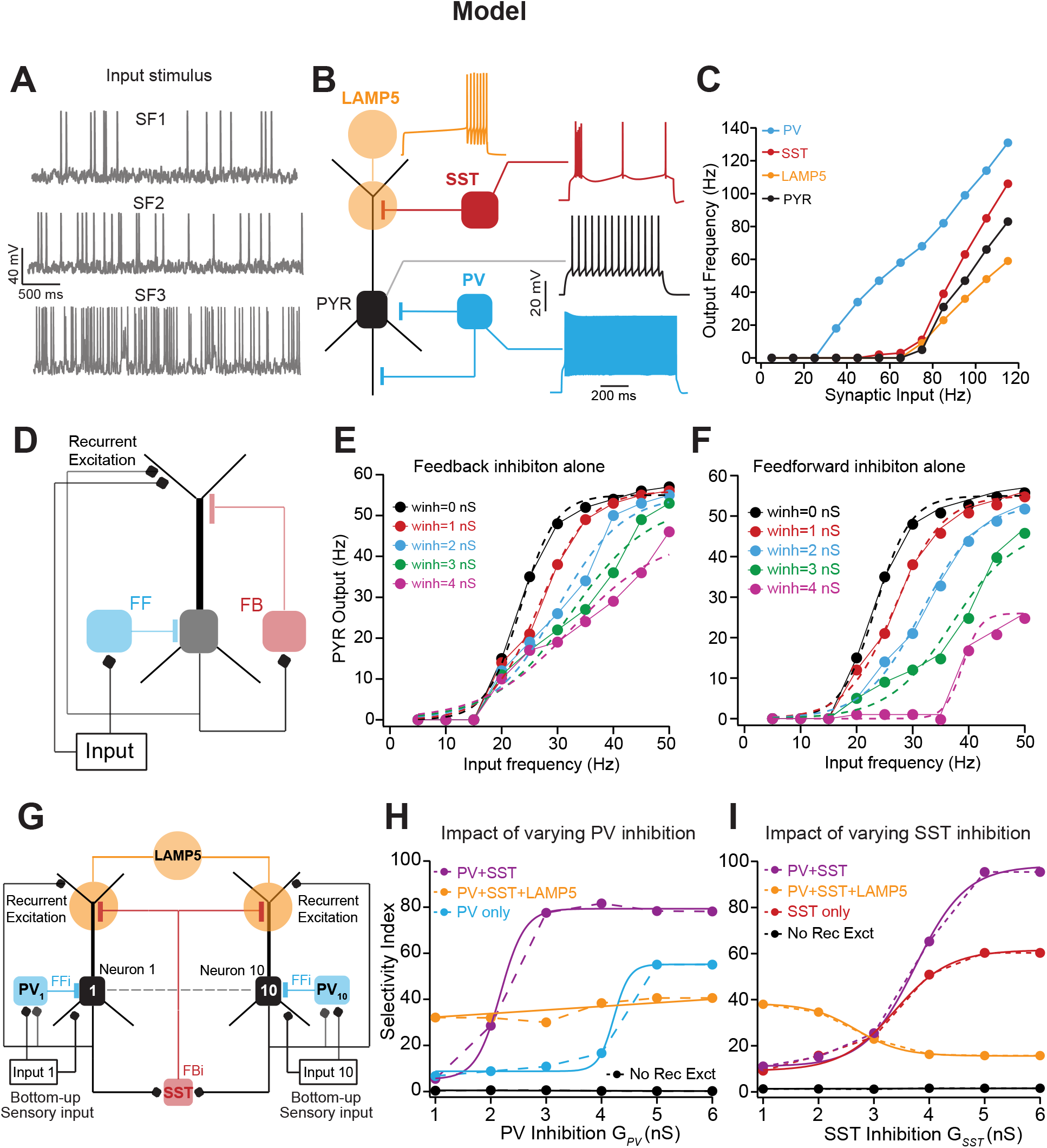
A) Voltage traces depicting neuronal firing for Poisson-distributed synaptic stimulation at various stimulus frequencies (SF) varying from 5 to 100 Hz. B) Schematic showing inhibitory neurons preferred synaptic location onto pyramidal neurons and their action potential firing dynamics of pyramidal neuron (black) and distinct interneuron types: PV (blue), SST (red), Lamp5 (orange). C) Plot showing output firing frequency (FF) as a function of SF for individual neuron types as described in (B). D) Schematic showing simplified circuit organization of feedforward (FF) and feedback (FB) inhibition on pyramidal neurons. E-F) Quantification of Feedforward (E) and Feedback (F) inhibition impact on the pyramidal neuron’s input-output function for different inhibitory synaptic conductance shown in various colors. Input to the network is Poisson-distributed synaptic stimulation at various SF as described in (A). G) Schematic showing the canonical cortical microcircuit motif. H-I) Impact of varying PV (H) and SST (I) inhibition on the selectivity index, which is calculated by subtracting the responses of pyramidal neurons to closely tuned inputs and multiplying the response difference with network gain. Selectivity index quantification as a function of inhibitory conductance strength for various scenarios such as in the presence of both PV, SST, and Lamp5 inhibition (orange), PV and SST only (magenta), PV only (cyan), SST only (red) and no recurrent excitation (black).

**Supplementary Figure 3:**
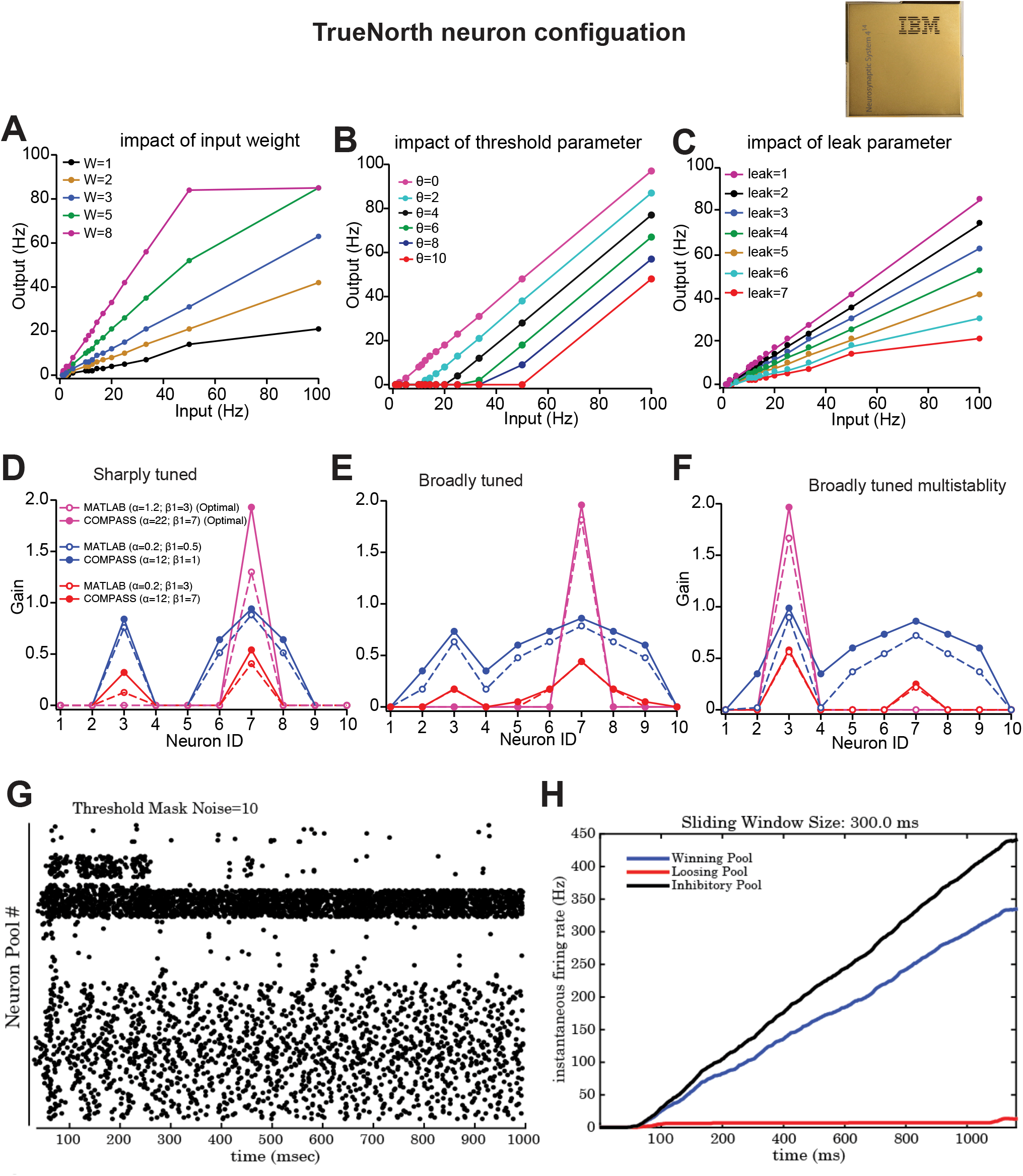
A-C) Impact of excitatory synaptic weight (A), threshold parameter (B), and leak parameter (C) on TrueNorth neuron input-output function for various magnitudes of the tested parameter shown in the corresponding color. D-F) Comparison of network gain of the rate-based neural network model implemented in the programming platform and with the TN neural networks implemented in the Compass (TrueNorth’s) emulator. For sharply tuned (D), broadly input with noise (E) and broadly tuned input with multistability (F) under optimal (magenta) and suboptimal (blue) parameter tuning. G-H) Impact of threshold mask noise parameter on WTA dynamics in TN. Spike raster plot of winning population showing WTA behavior for a threshold mask noise value of 10 (G). Population response dynamics of the winning and losing excitatory pool along with inhibitory population (H).

**Supplementary Figure 4:**
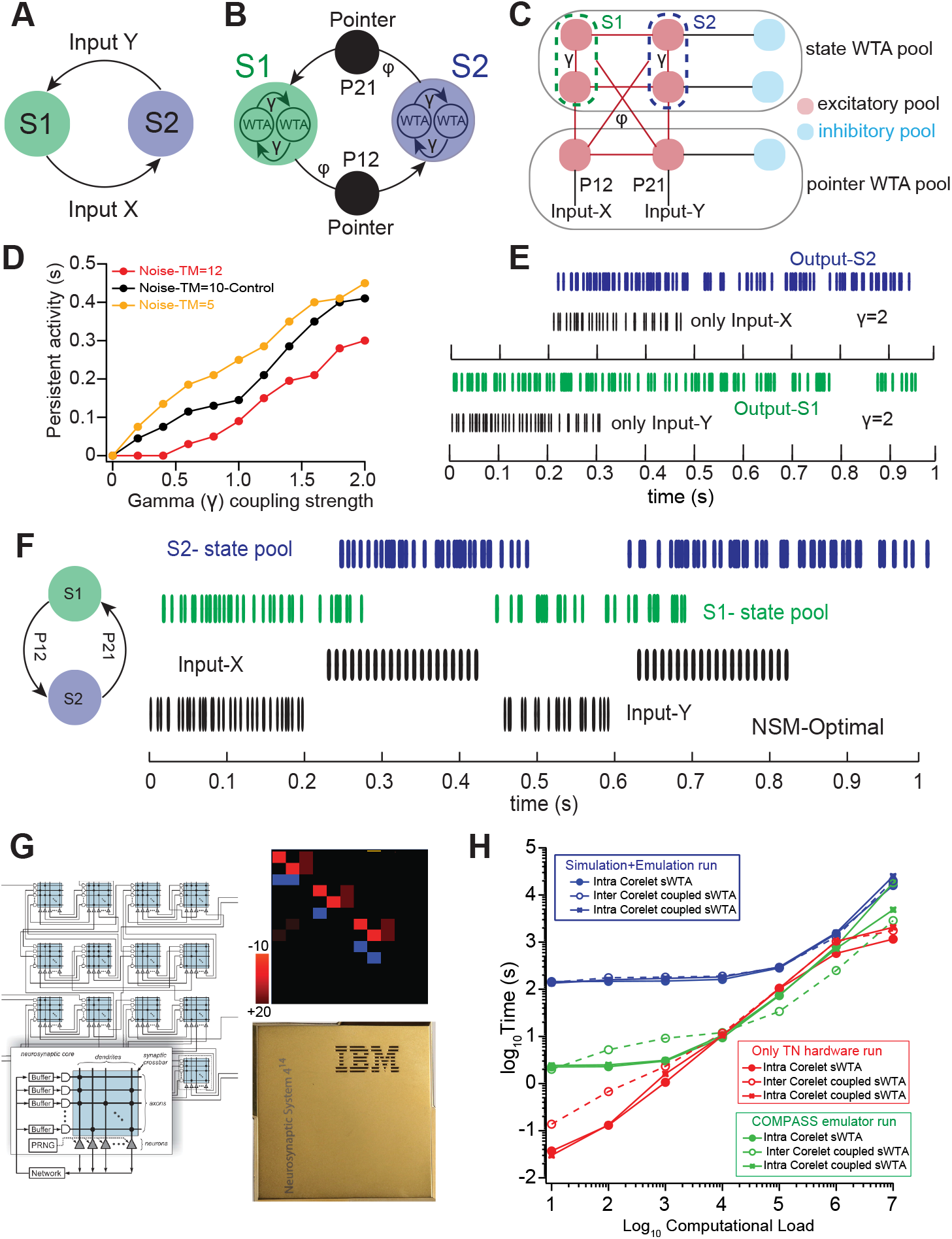
Neural state machines implemented with coupled sWTA motifs. A-B) Input-dependent state transition conditions in a two-state (S1 and S2) finite state automaton (FSA) and its neuronal implementation via coupled sWTA through excitatory connection, gamma (γ). The state transition is mediated through state pointer neurons. C) Population-level implementation of the two-state FSA as coupled sWTA in spiking-mode configuration for the two states (S1 and S2) and state-transition pointer (P12, P21) sWTA population as shown in (B) showing excitatory (red) and inhibitory (blue) neurons along with state pointer neurons (black) and state transition inputs. D) The impact of coupling strength () between sWTA and threshold mask noise on the persistence of the state maintenance of various threshold mask noise (TM) is shown in different colors. E) Demonstration of persistent activity for state maintenance with optimal parameters in the presence of only input X or input Y. S2 output in blue and S1 output in green. F) Raster plot of the state pool neurons showing state transitions from S1 (green) to S2 (blue) along with inputs for optimal parameters. G) Heatmap of the synaptic weight matrix implementing population-level two-state FSA in TN neurons. H) Hardware load analysis for various core size activation runtime for coupled sWTA network and comparing it with Truenorth versus software emulation and programming platform implementation. The log10 plot shows the differences in execution time as a function of computational load for hardware vs simulation. The graph shows the execution time for 1)NSCS (only TN Compass simulator) in green. 2) Native TN (only hardware) in red. 3) Total emaulation+simulation run time in blue. For each case, various configuration is explored with coupled sWTA implemented using within core neurons (intra-core) vs external core neurons (inter-core).

**Supplementary Figure 5:**
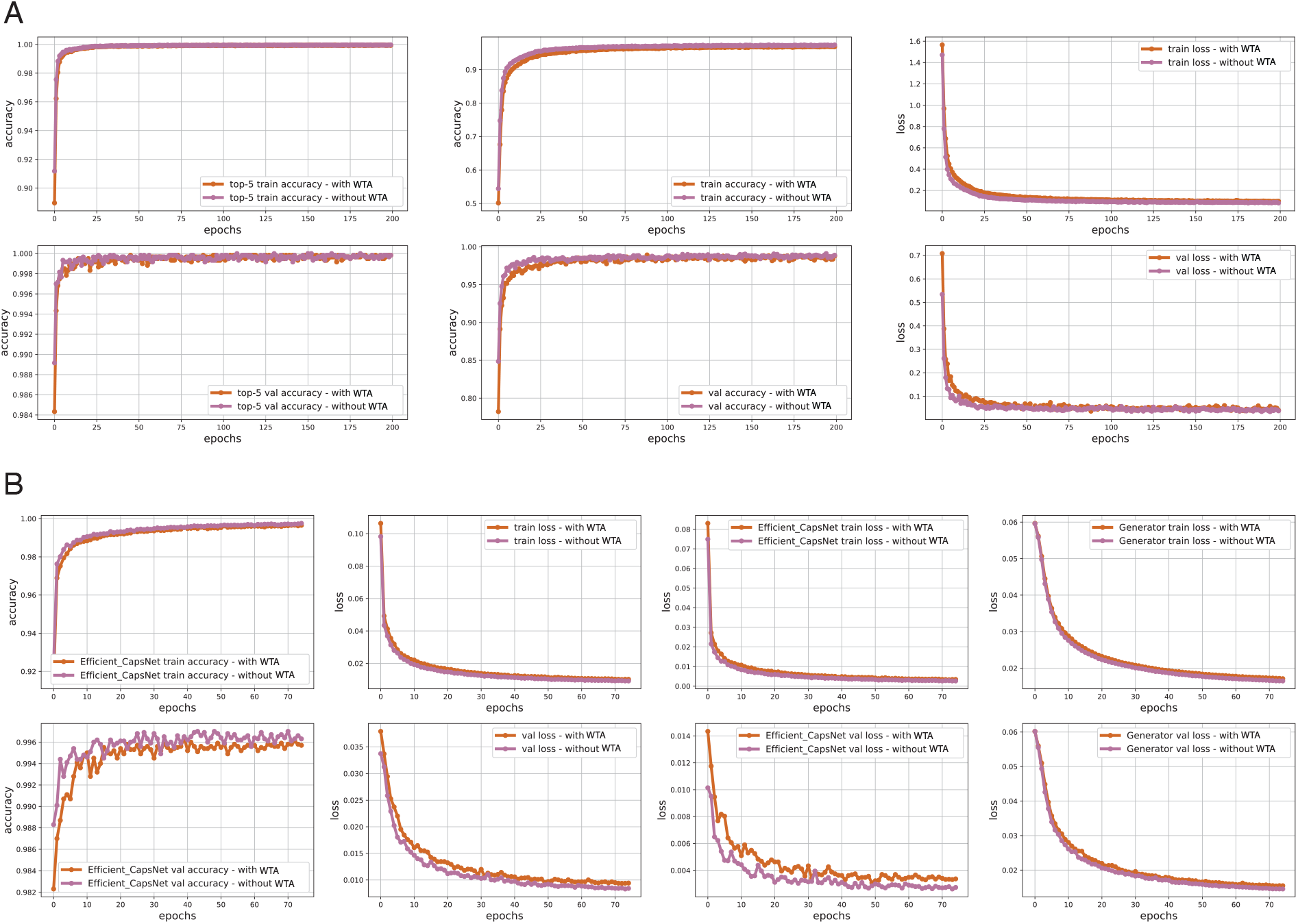
A. Training and validation performance (top-5 accuracy, total accuracy, and loss) of ViT architecture with and without adding WTA-layer. B. Training and validation performance (accuracies and losses) of CapsuleNet architecture with and without adding WTA-layer.

**Supplementary Figure 6:**
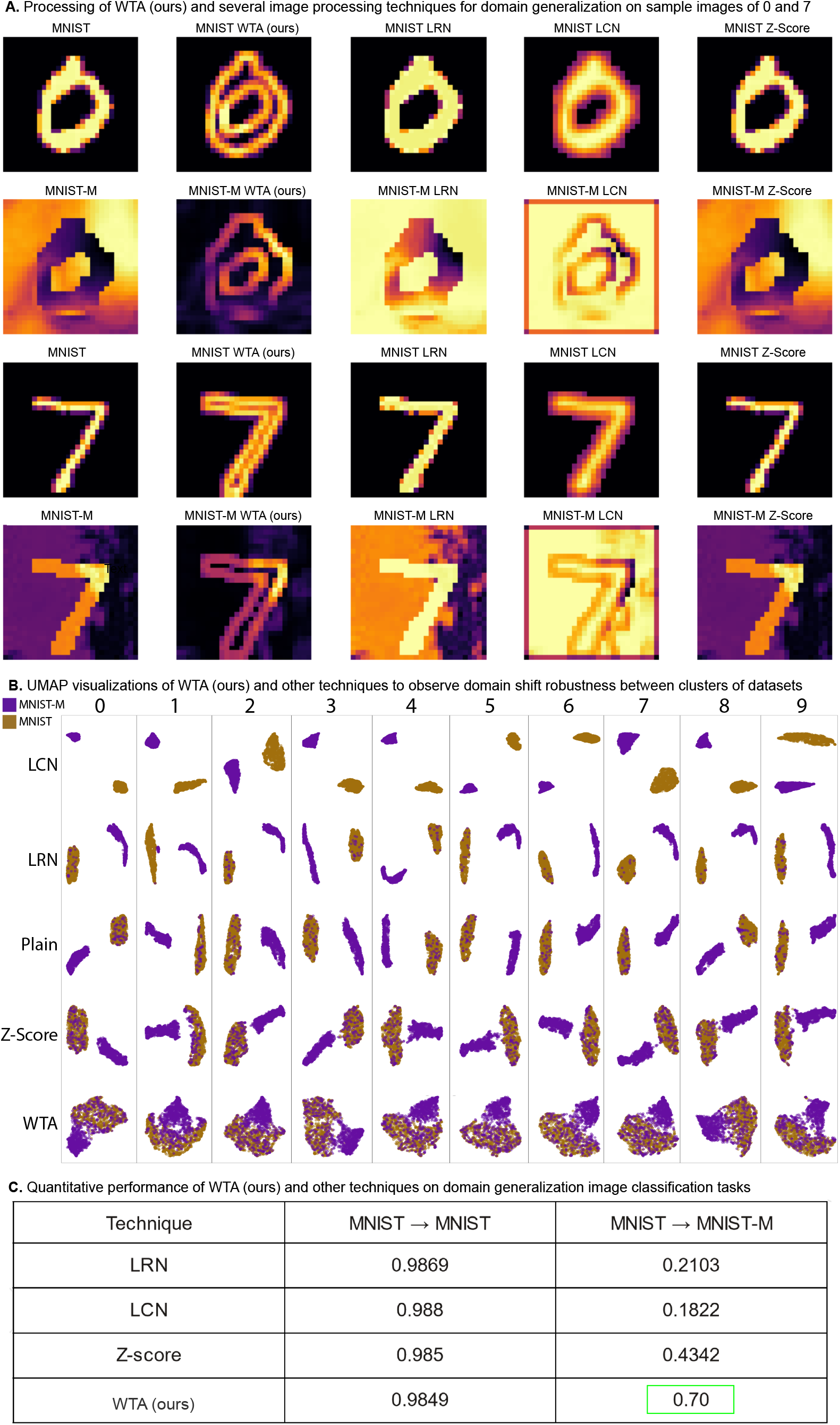
(A). Comparison of WTA and other normalization (LCN, LRN, Z-Score) techniques on 0 and 7 sample images from MNIST and MNIST-M. (B). UMAP embeddings of MNIST (brown) and MNIST-MM (purple) datasets, each row shows the embeddings of digits (0: leftmost - 9: rightmost) drawn from MNIST and MNIST-MM datasets for different techniques. (C). ViT is trained on MNIST and tested on both MNIST and MNIST-M datasets for comparisons between WTA and other techniques.

**Supplementary Figure 7:**
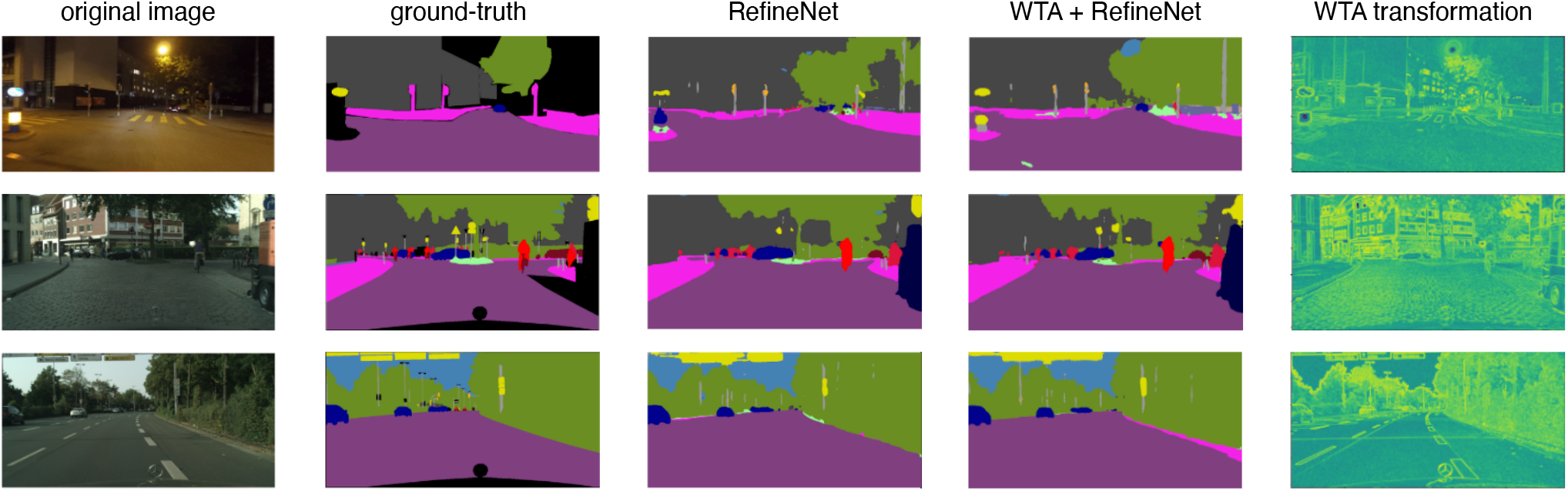
Qualitative outputs of Night-time driving data set against the RefineNet trained with WTA and without WTA layer. The WTA representation is demonstrated in the last column which is fed to RefineNet for training.

**Supplementary Table 1:** Performance comparison of various DNN architectures after and before adding WTA-layer. **Bold green** shows the benchmarking results, whereas normal green shows the performance improvement after adding the WTA layer but no benchmarking. The gray shows the cases where model performance is not improved by adding a WTA layer in the architecture (best seen in color).

| Models | MNIST<br>→<br>USPS<br>(%) | USPS<br>→<br>MNIST<br>(%) | SVHN<br>→<br>MNIST<br>(%) | MNIST<br>→<br>MNIST-M<br>(%) | MNIST-M<br>→<br>MNIST<br>(%) |
| --- | --- | --- | --- | --- | --- |
| Vision Transformer | **84.26 / 75.2**<br>*k* = [3,3] | **78.0 / 72.0**<br>*k* = [3,3] | **73.6 / 66.6**<br>*k* = [3,3] | 70.0 / 42.0<br>*k* = [3,3] | 98.1 / 98.3<br>*k* = [3,3] |
| EfficientNet | **83.5 / 77.9**<br>*k* = [4,4] | 74.2 / 50.7<br>*k* = [4,4] | 69.1 / 61.0<br>*k* = [3,3] | 48.8 / 18.8<br>*k* = [4,4] | **96.3 / 95.0**<br>*k* = [3,3] |
| CapsuleNet | 94.1 / 96.4<br>*k* = [3,3] | **87.8 / 87.2**<br>*k* = [3,3] | **75.9 / 58.1**<br>*k* = [4,4] | 57.2 / 22.0<br>*k* = [4,4] | **98.6 / 98.48**<br>*k* = [4,4] |
| MobileNet | 82.5 / 84.4<br>*k* = [4,4] | 70.9 / 60.0<br>*k* = [4,4] | **73.5 / 72.2**<br>*k* = [4,4] | **73.4 / 33.9**<br>*k* = [4,4] | **97.8 / 97.3**<br>*k* = [3,3] |
| ResNet | **82.8 / 82.5**<br>*k* = [3,3] | 66.0 / 58.5<br>*k* = [3,3] | **71.2 / 63.4**<br>*k* = [3,3] | 70.2 / 38.2<br>*k* = [3,3] | **97.8 / 97.4**<br>*k* = [3,3] |
|  | *previous best:*<br>82.2 | *previous best:*<br>77.5 | *previous best:*<br>70.58 | *previous best:*<br>70.28 | *previous best:*<br>97.8 |

**Supplementary Table 2:** Training settings for each model for object classification and segmentation tasks.

| Architecture | Optimizer | Learning Rate | Loss Function | Epochs<br>(saving best model by validation loss) |
| --- | --- | --- | --- | --- |
| Transformer | Adam | 0.001 | crossentropy | 200 |
| CapsuleNet | Adam | 0.0005 | margin loss + 'MSE' | 75 |
| EfficientNet | Adam | 0.001 | crossentropy | 100 |
| MobileNet | Adam | 0.01 | crossentropy | 100 |
| ResNet | Adam | 0.01 | crossentropy | 100 |
| U-Net | Adam | 0.0001 | DICE-based loss | 100 |
| Refine Net | SGD | Decaying | crossentropy | 120 |

